# Temporal stability for individual differences in both habit learning and expression

**DOI:** 10.64898/2026.11.04.764200

**Authors:** Pablo Martínez-López, José J. León, Francisco Garre-Frutos, Antonio Vázquez-Millán, Sara Molinero, Tamara Giménez-Fernández, Miguel A. Vadillo, David Luque

## Abstract

Actions transition from goal-directed to stimulus-driven habits through repetition. Individual differences in the capacity to learn and control habits are central to current models of psychopathology, which assume that overreliance on habitual control drives maladaptive behaviors. However, whether these tendencies reflect stable, trait-like cognitive processes remains unclear. We examined the temporal stability of individual differences in habit learning and expression. Across two preregistered experiments (total N = 313), we assessed stimulus–response habits using the same stimuli after one day or different stimuli after four months. Evidence for stable individual differences in habit commissions was strong and comparable across experiments, with computational modelling further supporting trait-like tendencies for habit learning and expression. Trait-like tendencies were also reflected in action automatization while practicing stimulus-response associations, which was associated with future habitual behavior. Our findings have theoretical implications for our understanding of habit formation and its role in psychopathology.

---

There are occasions in which people seem to have no control over their own actions. One may try to limit snacking between meals^1,2^ or social media use before going to sleep^3,4^, often unsuccessfully. Neurocognitive theory posits that repetition transforms deliberate actions into habits performed rapidly and with little conscious deliberation^5,6^. While habit learning facilitates adherence to healthy routines^7,8^, maladaptive perseverant behaviors are often conceptualized as habits resistant to change^9,10^. Prominent psychiatric models further propose that an inherent predisposition to overrely on habitual control increases vulnerability to compulsive symptoms^11–13^.

Experimentally, habits are conceptualized as learned stimulus–response associations that persist even when goals change. Standard laboratory protocols evaluate habit formation by training an action over several days and testing participants’ ability to adapt when that learned response becomes counterproductive^14–16^. Substantial effort has focused on linking experimental habits to psychopathology^17^, under the key theoretical assumption that individual differences in habitual control reflect stable, trait-like tendencies. However, previous studies have assessed experimental habits at a single time point immediately following training^14,18^. Consequently, it remains unknown whether interindividual variation in habitual behavior reflects temporally stable cognitive processes or transient states. Because habitual control relies on both the rate of habit acquisition and the degree of habit expression, demonstrating the temporal stability of both components is essential to establish habit formation as a cognitive trait underlying long-term behavioral repertoires.

Across two preregistered experiments, we examined the temporal stability of individual differences in habitual behavior using test-retest designs. Habitual behavior was assessed using either the same overtrained set of stimulus–response associations (Exp. 1, *N* = 79) or a different set of overtrained associations at each test (Exp. 2, *N* = 170), with a short interval between tests in Exp. 1 (∼24 hours) and a longer interval in Exp. 2 (∼4 months).

In both experiments, participants learned sets of four stimulus–response associations. To induce habit formation, they practiced an initial stimulus–response mapping (1,000 trials/day; Exp. 1, 3 days; Exp. 2, 4 days; Fig. 1a, b). Previous work has shown that when a well-trained stimulus–response association changes, whether individuals produce the habitual or goal-directed response depends on the time available for response preparation^13,19–23^. Indeed, testing participants under varying levels of time pressure appears necessary to reveal the extent to which a repeated action has become habitual^14^. To uncover latent habit formation of the initially practiced responses, participants completed forced-response test trials in which the time available to prepare a response varied from 0 ms to 1,200 ms across trials (Fig. 1c)^19^. Previous results indicate that responses are predominantly random when the preparation time is too short to process the stimulus (< ∼300 ms). Following previous work^19,20,22^, we quantified habitual responding (i.e., pressing the original key for remapped associations) within a participant-specific preparation-time window from *t*_min_, the minimum preparation time at which responses to consistent associations become nonrandom (Fig. 1d; ∼300 ms of preparation time, see Supplementary Fig. 1), to *t*_min_ + 300 ms.

**Figure 1:**
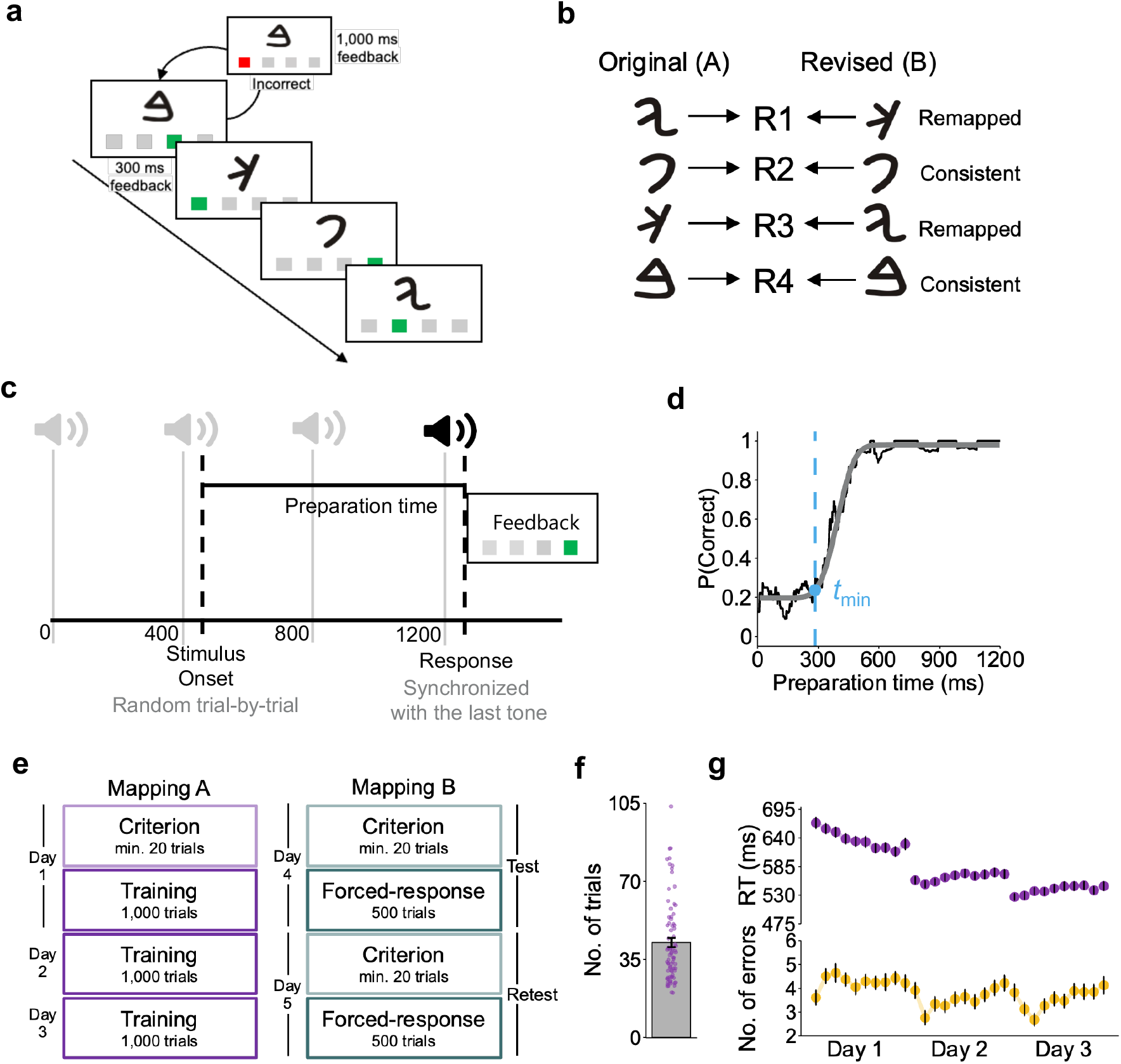
Experimental protocol for habit induction and testing. **(a)** Trial structure of the criterion and training blocks. Participants completed blocks of 100 trials (10 blocks per training session). A 1-s time penalty was applied for incorrect responses, and a correct response was required to end the trial. During the training blocks, participants were instructed to respond as quickly and accurately as possible. During the criterion blocks, they received no time-pressure instructions and were asked to learn the associations through trial and error. **(b)** Example of the stimulus–response associations learned by participants. **(c)** Schematic of the forced-response method. On each trial, participants heard a four-tone metronome sequence (400 ms intervals) and were required to respond precisely on the final beat. To manipulate the time available for response preparation, the stimulus appeared at a randomly selected time between 0 and 1,200 ms before the fourth tone, sampled from a uniform distribution. Preparation time was calculated as the interval between stimulus onset and the participant’s response. In the example shown, the trial began, the stimulus appeared at 450 ms, and the participant responded at 1,250 ms (50 ms after the final beat), yielding a preparation time of 800 ms. **(d)** Representative data illustrating the expected relationship between preparation time and the probability of a correct response on forced-response trials for consistent associations. A cumulative Gaussian function was fitted (gray line) to the observed data (black line; proportion of correct responses within a 100-ms sliding window) with *t*_min_ denoting the preparation time at which the fitted function reached 5% of its height. **(e)** Overview of the Exp. 1 design. All participants followed the same procedure: they learned and extensively practiced four stimulus–response associations (Mapping A). To assess habit formation, the initial mapping was revised (Mapping B), and the responses associated with two stimuli were switched (see panel b). Participants were instructed to respond according to Mapping B during both the test and retest sessions. **(f)** Mean number of trials required to reach the learning criterion for the original mapping (Mapping A) in Exp. 1, defined as five consecutive correct responses for each stimulus. Each dot represents one participant, and the error bar indicates the standard error. **(g)** Data from Exp. 1 training stage. Each point represents one block of 100 trials. Mean response times (RTs) for correct responses are shown in the upper panel, and error rates are shown in the lower panel. Error bars indicate standard errors.

## Results

### Short-term stability of individual differences in habit expression

Performance on cognitive tasks following learning is often used to assess the extent to which a behavior has been acquired. However, individual differences in performance—that is, true variability in the ability to respond in a goal-directed versus habitual manner— necessarily affects inferences about variability in the learning process itself, including individual differences in the rate of habit acquisition^24^. Accordingly, before examining the long-term temporal stability of habit formation, in Exp. 1 we sought to establish that individual differences in the expression of learned habits were not purely transient or context-dependent. We therefore analyzed habit expression across test–retest sessions separated by only 1 day (Fig. 1e), allowing us to assess whether the same stimulus–response habit was expressed consistently across tests conducted on consecutive days.

Before examining individual-level performance across tests, we assessed group-level changes, as these could otherwise obscure the consistency of individual performance over time^25,26^. In Exp. 1, participants first learned the stimulus–response associations (Mapping A) through trial-and-error in a criterion block that required five consecutive correct responses for each association (42.54 ± 2.07 trials; mean ± s.e.m.; Fig. 1f). They then practiced the associations for 3 days. Performance improved across training, as reflected in reduced reaction times (RTs) for correct responses (Fig. 1g), whereas error rates remained stable. This pattern indicates that the initial learning criterion was sufficient to ensure reliable acquisition of stimulus–response associations.

One day after the last training session, participants completed a criterion block in which they learned a revised stimulus–response mapping (Mapping B). Two associations were remapped, whereas the other two remained consistent (Fig. 1b, Fig. 2a). The same Mapping B criterion block was completed at the beginning of the retest session on the following day. Participants required fewer trials to meet the learning criterion on the retest day (difference, 14.29 ± 6.44 trials; *t*_78_ = 2.21; *P* = .029; *d*_z_ = 0.25, 95% confidence interval (*CI*) [0.02, 0.48]; Fig. 2b), indicating that they retained the Mapping B associations learned the previous day. Forced-response trials revealed time-dependent habit expression in both test and retest: participants produced the correct remapped responses at long preparation times (>900 ms) but were biased toward the original habitual Mapping A at shorter latencies (300–600 ms). Habitual responding decreased at retest (Fig. 2c), although habitual errors remained more frequent than responses that had never been associated with the stimulus (other errors) in both sessions (test: difference, .10 ± .01; preregistered one-tailed *t*_78_ = 6.97; *P* < .001; *d*_z_ = 0.79, 95% *CI* [0.53, 1.04]; retest: difference, .02 ± .01; preregistered one-tailed *t*_78_ = 2.20; *P* = .015; *d*_z_ = 0.25, 95% *CI* [0.02, 0.47]; Fig. 2d). Because other errors remained stable (difference, .01 ± .007; *t*_78_ = 1.64; *P* = .104; *d*_z_ = 0.19, 95% *CI* [-0.04, 0.41]) while habitual errors decreased (difference, .08 ± .01; *t*_78_ = 8.19; *P* < .001; *d*_z_ = 0.93, 95% *CI* [0.66, 1.19]), these findings indicate that, although habits persisted, retesting facilitated retrieval of the goal-directed responses and reduced habitual behavior.

**Figure 2:**
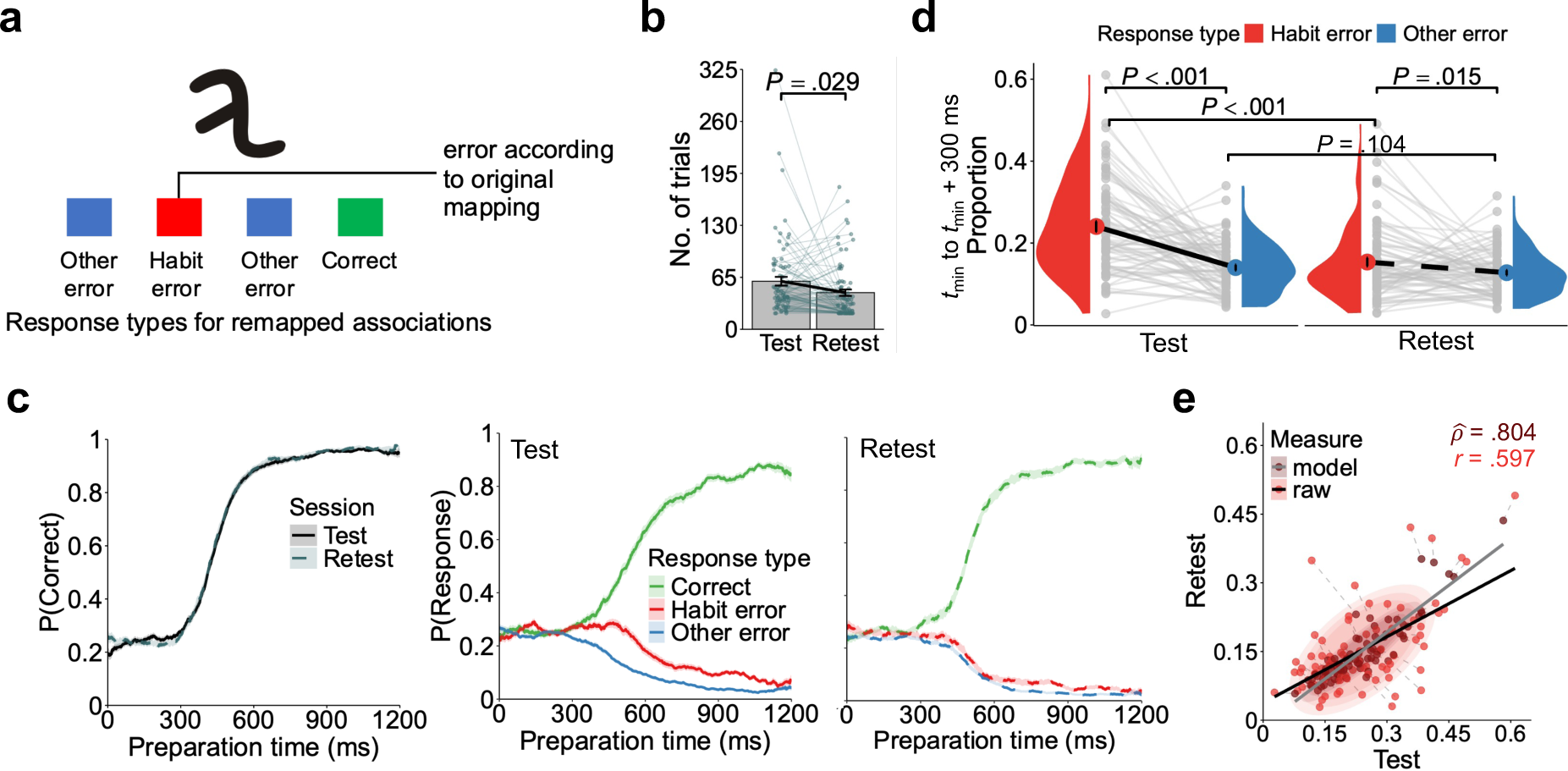
Consistency of habit expression across tests and individuals. **(a)** Response types for associations remapped under Mapping B. Trials were classified according to whether participants produced the new correct response (response switch), a habitual error (previously practiced response), or another error (a response never previously associated with the stimulus). **(b)** Mean number of trials required to reach the learning criterion for the revised Mapping B. Each dot represents one participant, and error bars indicate the standard error. The *P*-value corresponds to a two-sided *t*-test. **(c)** Performance on the forced-response trials as a function of preparation time. From left to right, 100-ms sliding windows show the probability of correct responses for consistent associations (first panel), the probability of each response type for remapped associations during the test session (second panel), and the corresponding probabilities during the retest session (third panel). Shaded areas indicate standard errors. **(d)** Data are presented as half-violin plots. Colored points indicate group means, error bars indicate standard errors, and gray points represent individual participants. Within-session *P*-values correspond to preregistered one-tailed *t*-tests, whereas between-session *P*-values correspond to two-sided *t*-tests. **(e)** Test–retest correlation in habit expression. Each red point represents a participant’s proportion of habitual errors from *t*_min_ to *t*_min_ + 300 ms based on the raw data. Darker points represent estimates from a Bayesian generative model used to correct the observed correlation for attenuation due to measurement error and show the means of the participant-level posterior distributions for the proportion of habitual errors. Dashed lines connect the raw and model-based estimates for each participant.

We then examined whether individual differences in habit expression remained stable across tests. Crucially, the proportion of habitual errors was strongly correlated between the test and retest sessions (preregistered one-tailed *r*_77_ = .59; *P* < .001; Fig. 2e), indicating stable individual differences in habitual behavior. Although our habit measure showed appropriate psychometric reliability across both test sessions (test: *r*_sb_ = .77, 95% *CI* [.69, .84]; retest: *r*_sb_ = .74, 95% *CI* [.65, .83]), the measurement error could still have attenuated the observed correlation^27,28^. To account for this measurement noise, we further characterized the observed correlation using Bayesian generative models^29,30^, which revealed a strong latent association after accounting for measurement error between sessions (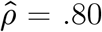, 95% credible interval (*CrI*) [.55, .93]; Fig. 2e). Notably, other errors were not associated with habitual errors in either session (all *r*_77_’s ≤ .03; *P*’s ≥ .750). Thus, individual differences in habit expression were stable after 1 day.

### Habit learning and expression follow trait-like tendencies

We next examined whether individual differences in habitual behavior remained stable over longer periods of time and when participants have to learn different contents. Demonstrating such stability requires that individual differences in both the rate of habit acquisition and subsequent habit expression remain stable across assessments. Short intervals between assessments may increase behavioral consistency by reducing the likelihood of transient effects^26,31^. However, trait-like psychological constructs should remain stable over longer time periods. Therefore, to examine trait-like tendencies in habitual behavior, in Exp. 2 we assessed two distinct sets of stimulus–response associations approximately 4 months apart (mean interval = 118.42 days, standard deviation (*SD*) = 7.51, range = 105–133; Fig. 3a).

**Figure 3:**
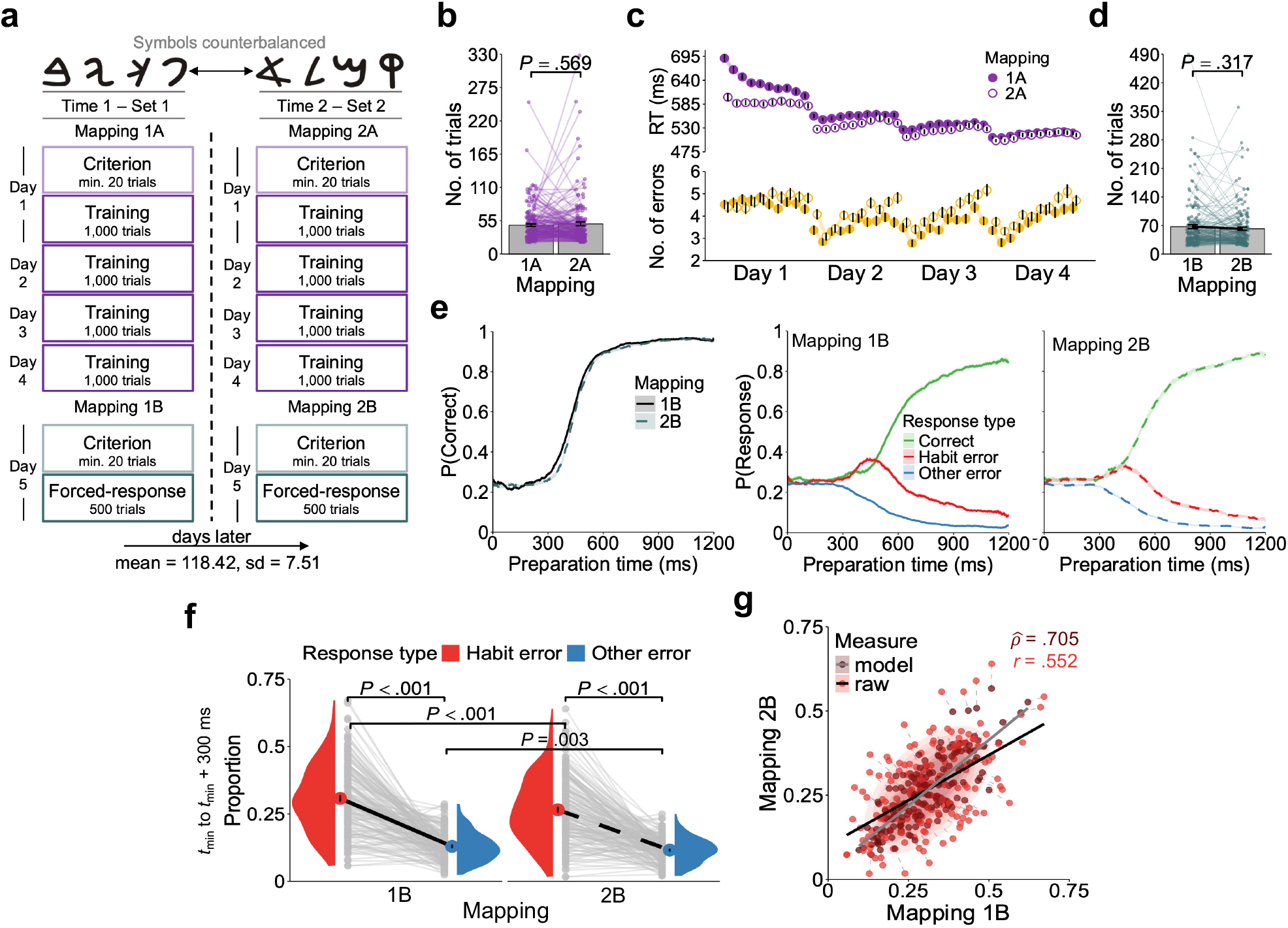
Long-term stability of individual differences in habitual behavior across distinct habits. **(a)** Overview of Exp. 2 design. All participants followed the same procedure: habit formation was induced and tested using one set of stimuli, and approximately 4 months later, participants repeated the procedure using a different stimulus set. Thus, both habit formation and expression had to remain stable to yield consistent individual differences across tests. Mappings were classified as original (Mapping A) or revised (Mapping B), and as involving the first (1) or second (2) stimulus set, yielding Mappings 1A, 1B, 2A and 2B. **(b)** Mean number of trials to reach the learning criterion for the extensively trained A mappings. Each dot represents a single participant, and error bars denote the standard error. The *P*-value corresponds to a two-sided *t*-test. **(c)** Data from the training stage. Each point represents one block. The upper panel shows mean RTs for correct responses, and the lower panel shows error rates. Error bars denote standard errors. **(d)** Data from the criterion blocks for each revised mapping (as in panel b). **(e)** Performance as a function of preparation time, showing responses for consistent associations across the original mappings (first panel), remapped associations under Mapping 1B (second panel), and remapped associations under Mapping 2B (third panel). **(f)** Group-level and individual data presented as half-violin plots (as in Fig. 2d). **(g)** Correlation in habitual behavior across stimulus sets over the 4-month interval. Red points represent the raw data, whereas darker points represent estimates from the generative model used to correct the observed correlation for attenuation due to measurement error.

In Exp. 2, the training phase was extended to four consecutive days. The two sets of stimulus–response associations trained (first training stage, Mapping 1A; second training stage, Mapping 2A) were matched in complexity, as the number of trials required to reach the learning criterion (five consecutive correct responses per association) did not differ significantly between training phases (difference, 2.05 ± 3.61 trials; *t*_169_ = 0.57; *P* = .569; *d*_z_ = 0.04, 95% *CI* [-0.11, 0.19]; Fig. 3b). Performance improved across both training phases, although RTs for the second set of stimuli were faster from the outset, likely reflecting familiarity with the experimental protocol (Fig. 3c).

During the test sessions, the number of trials required to learn the revised mappings, Mapping 1B in the first test session and Mapping 2B in the second, was similar between assessments (Fig. 3d; difference, 4.92 ± 4.91 trials; *t*_169_ = 1.00; *P* = .317; *d*_z_ = 0.08, 95% *CI* [-0.07, 0.23]). Forced-response trials revealed the same time-dependent pattern of habit expression for both mappings as observed in Exp. 1 (Fig. 3e), while adding an extra day of training increased habit errors when we compared the first test session of each experiment (difference, .06 ± .01; *t*_167.76_ = 4.20; *P* < .001; *d* = 0.65, 95% *CI* [0.34, 0.96]; Fig. 2d versus Fig. 3f). In both assessments of our second experiment, the proportion of habitual errors was significantly greater than that of other errors (Mapping 1B: difference, .17 ± .01; preregistered one-tailed *t*_169_ = 16.40; *P* < .001; *d*_z_ = 1.26, 95% *CI* [1.06, 1.46]; Mapping 2B: difference, .14 ± .009; preregistered one-tailed *t*_169_ = 15.31; *P* < .001; *d*_z_ = 1.18, 95% *CI* [0.98, 1.37]; Fig. 3f). Unexpectedly, both habitual and other errors were significantly less frequent for Mapping 2B than for 1B (habit error: difference, .04 ± .009; *t*_169_ = 4.65; *P* < .001; *d*_z_ = 0.36, 95% *CI* [0.20, 0.51]; other errors: difference, .01 ± .004; *t*_169_ = 2.95; *P* = .003; *d*_z_ = 0.23, 95% *CI* [0.07, 0.38]; Fig. 3f). Thus, a general practice effect was evident at the second assessment: goal-directed control improved across assessments despite the use of different stimulus–response associations.

Crucially, the proportion of habitual errors was strongly correlated between tests (preregistered one-tailed *r*_168_ = .55; *P* < .001; Fig. 3g), despite the four-month interval and the group-level practice effects. After accounting for measurement error (Mapping 1B: *r*_sb_ = .77, 95% *CI* [.72, .82]; Mapping 2B: *r*_sb_ = .78, 95% *CI* [.73, .83]), the estimated association was even stronger 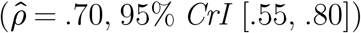. Other errors were not associated with habitual errors at either test (all *r*_168_’s ≤ .08; all *P*’s ≥ .293). Notably, the magnitude of this association was similar to that observed in Exp. 1, in which habitual behavior was assessed using the same associations and the test and retest sessions were separated by only 1 day. Additionally, we conducted a multiverse analysis, which showed that the observed temporal stability was robust across 96 a priori valid pipelines (Supplementary Fig. 2). These findings provide strong evidence that individual differences in habit learning and expression are temporally stable, suggesting that they reflect an enduring behavioral phenotype rather than transient states.

### Computational modelling of habitual behavior

Using a previously established computational framework for forced-response data^19,22,32^, we aimed to further characterize group- and participant-level data in each experiment. We pitted our data against an all-or-nothing model, a continuum model, and a no-habit model (Fig. 4a and *Methods*). The all-or-nothing model assumes habitual and goal-directed responses are prepared at random times following two independent cumulative Gaussian distributions and predicts habit behavior based mainly on the degree of overlap between them (i.e., speed difference, µ_Δ_). Participants would express habitual responses depending on whether, for the available time in a given forced-response trial, the habitual response was prepared while the goal-directed response was not. The all-or-nothing model thus would emit a habitual response whenever it is prepared unless overridden by goal-directed control. The continuum model incorporates a habit strength parameter (0 ≤ W_H_ ≤ 1) that determines the proportion of trials in which the habitual response is prepared. While in the all-or-nothing model the habitual response is eventually prepared on every trial (W_H_ = 1), in the continuum model the probability of preparing a habitual response is modulated by habit strength, with stronger habit strength increasing the likelihood that the habitual response is available, thereby leading to habit errors more frequently. The no-habit model assumes responses are governed only by goal-directed control.

**Figure 4:**
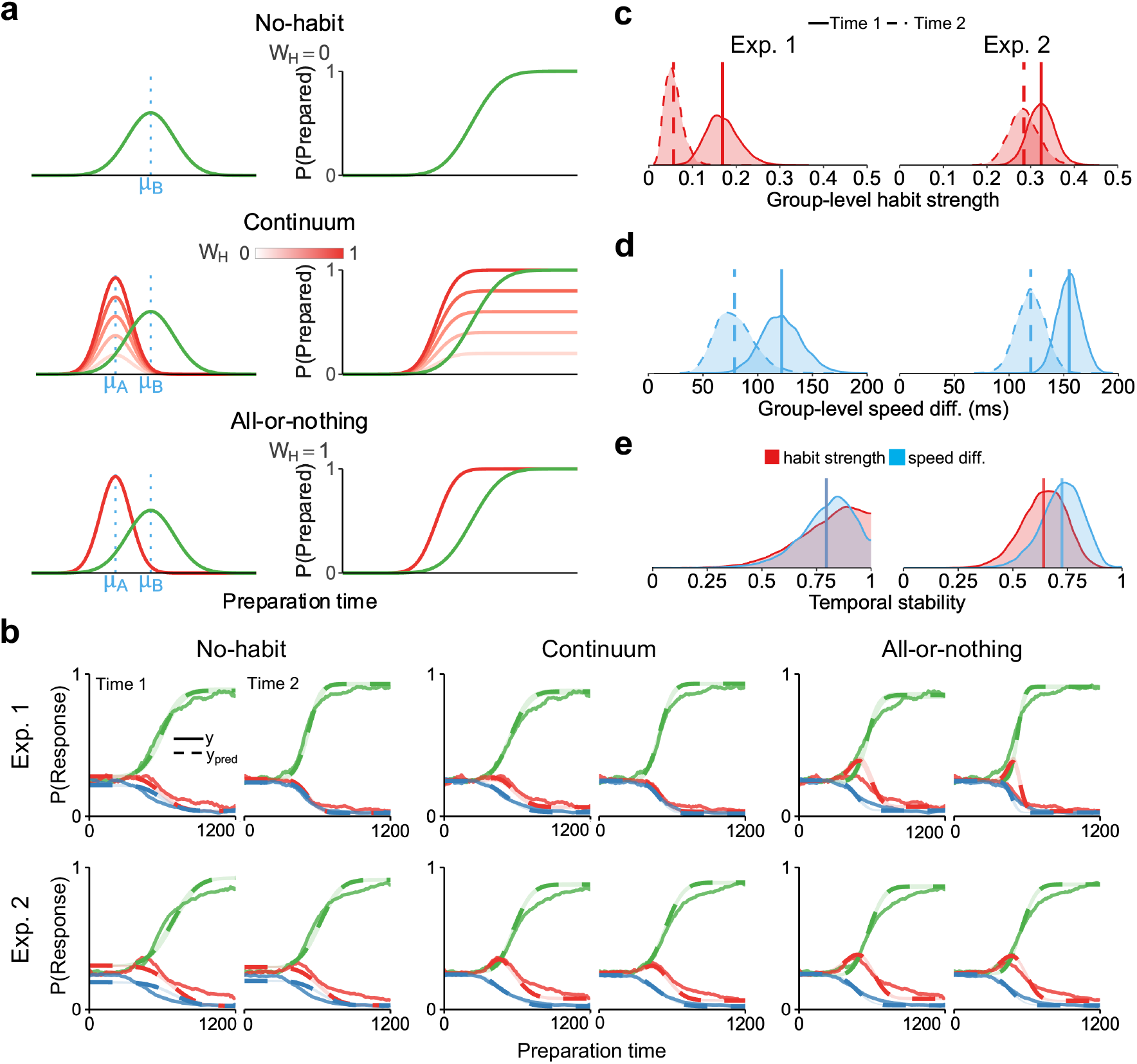
Comparison of computational models, dynamic and temporal stability of the main parameters. **(a)** The no-habit model hypothesizes that the goal-directed response is prepared at a random time following a cumulative distribution function. Before the response is prepared, accuracy is expected to be at the chance level. The continuum model assumes that the habitual response is, on average, prepared earlier than the goal-directed one and follows a different cumulative distribution function. The height of that function is defined by the habit strength parameter (W_H_) and modulates the probability of preparing the habitual response. Once the goal-directed response is prepared, it replaces the habitual one. Therefore, habitual behavior is also modulated by the degree of overlap between the habitual and the goal-directed distributions. The all-or-nothing model fixes habit strength at its maximum (W_H_ = 1); hence, habitual behavior depends solely on the timing of each response. **(b)** Posterior predictive checks supported the continuum model best fit the observed data in both experiments. Time 1 and Time 2 refers to the first (Exp. 1, test; Exp. 2, Mapping 1B) and second assessment (Exp. 1, retest; Exp. 2, Mapping 2B) of each experiment. **(c)** Posterior distributions of the habit strength parameter across sessions and experiments. Lines denote the mean of each posterior distribution. **(d)** Posterior distributions of the speed diff. parameter across sessions and experiments. Lines reflect the mean of each posterior distribution. **(e)** Posterior distributions of the model-based correlation of habit strength and speed difference parameters in each experiment. Lines denote the mean of each distribution

For each experiment, we fitted a Bayesian hierarchical version of each model to remapped stimuli of both test sessions (see Supplementary Fig. 3 for prior predictive checks). We compared the three models using Pareto-smoothed importance sampling cross-validation (PSIS-LOO)^33^ and, as reported in Table 1, the continuum model achieved the highest predictive accuracy in both experiments in the joint fit and in each test session indexed separately (see Supplementary Fig. 4 for model recovery based on ELPD > 2 × SE). The continuum model also showed better descriptive adequacy than the other models, as illustrated by the posterior predictive checks (Fig. 4b). Therefore, we concluded that the continuum model, which included a habit strength parameter, better accounted for our data than the all-or-nothing and no-habit models.

**Table 1:** PSIS-LOO model comparison. ΔELPD refers to the difference between a model’s estimated expected log pointwise predictive density and that of the top-ranked model (Continuum model). ΔSE is the standard error of that difference. Time 1 and Time 2 refers to the first (Exp. 1, test; Exp. 2, Mapping 1B) and second assessment (Exp. 1, retest; Exp. 2, Mapping 2B) of each experiment. Global estimates reflect the joint fit of observations at Time 1 and Time 2, while the Time 1 and Time 2 columns show the ELPD contributions from their respective sessions. We conclude that the continuum model provides a better fit for each dataset, given that the differences in ELPD between each model and the continuum model are greater than 2 × SE^33^.

|  | Global |  | Time 1 |  | Time 2 |  |
|---|---|---|---|---|---|---|
| Model | ΔELPD | ΔSE | ΔELPD | ΔSE | ΔELPD | ΔSE |
| *Experiment 1* |  |  |  |  |  |  |
| Continuum | 0 | 0 | 0 | 0 | 0 | 0 |
| All-or-nothing | -371.2 | 27.0 | -157.5 | 17.6 | -213.7 | 20.5 |
| No-habit | -379.3 | 28.9 | -273.4 | 23.8 | -105.9 | 16.3 |
| *Experiment 2* |  |  |  |  |  |  |
| Continuum | 0 | 0 | 0 | 0 | 0 | 0 |
| All-or-nothing | -416.9 | 27.7 | -201.0 | 18.7 | -215.9 | 20.5 |
| No-habit | -1956.6 | 64.2 | -1132.3 | 48.9 | -824.3 | 41.7 |

### Group-level changes and temporal stability of habit strength

Then, we tested whether the habit strength parameter aligned with the group- and individual-level behavioral results previously reported in the article (see Supplementary Fig. 5 for parameter recovery using the design of both experiments). Habit strength increased after training the associations an extra day, as the difference between the first session of each experiment was positive for nearly all posterior samples (Exp. 1, posterior mean (*M*) = .16, 95% *CrI* [.10, .25]; Exp. 2, *M* = .32, 95% *CrI* [.26, .38]; difference, *M* = .15, 95% *CrI* [.05, .24], probability of direction (*pd*) = .998; Fig. 4c). There was evidence that habit strength was reduced in Exp. 1 after retesting the same associations (retest, *M* = .05, 95% *CrI* [.02, .10]; difference, *M* = .11, 95% *CrI* [.03, .19], *pd* = .998). In Exp. 2, the evidence for a change across tests, in which two different sets of 4-day trained associations were assessed, was uncertain (Mapping 2B, *M* = .28, 95% *CrI* [.22, .35]; difference, *M* = .03, 95% *CrI* [-.03, .11], *pd* = .848).

In both experiments, temporal stability of individual differences in the habit strength parameter closely resembled the effect sizes previously reported for the raw scores. Temporal stability correlations were estimated within the hierarchical model, as model-based correlations tend to provide more accurate estimates of the latent relationships than traditional methods^34–36^. As seen in Fig. 4e, in Exp. 1, the habit strength parameter showed a posterior mean test-retest correlation of 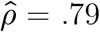, 95% *CrI* [.45, .99]. In Exp. 2, the temporal stability of the parameter between the test involving Mapping 1B and the test conducted four months apart involving Mapping 2B was 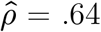, 95% *CrI* [.42, .82].

### Group-level changes and temporal stability of response speed

We also examined group-level changes and temporal stability in the difference (µ_Δ_) between the onset of the habitual (µ_*A*_) and goal-directed processes (µ_*B*_) (see Supplementary Fig. 5 for parameter recovery).

The evidence that the speed difference parameter changed after adding an extra day of training was uncertain (µ_Δ_; first test session of Exp. 1, *M* = 122 ms, 95% *CrI* [86, 161]; first test session of Exp. 2, *M* = 155 ms, 95% *CrI* [134, 177], difference, *M* = 33 ms, 95% *CrI* [-11, 75], *pd* = .934; Fig. 4d). In Exp. 1, not only was habit strength reduced after retesting, but also the speed difference parameter was possibly affected (retest, *M* = 78 ms, 95% *CrI* [48, 118]; difference, *M* = 43 ms, 95% *CrI* [-6, 90], *pd* = .956). In the retest session, participants were able to select the goal-directed response with less preparation time (µ_*B*_; test, *M* = 538 ms, 95% *CrI* [518, 558]; retest, *M* = 496 ms, 95% *CrI* [481, 511]; difference, *M* = 42 ms, 95% *CrI* [25, 59], *pd* = 1), while the evidence for a change in the onset of the habitual process was uncertain (µ_*A*_; test, *M* = 416 ms, 95% *CrI* [381, 449]; retest, *M* = 417 ms, 95% *CrI* [379, 448]; difference, *M* = -0.69 ms, 95% *CrI* [-44, 45], *pd* = .524).

In Exp. 2, the speed difference parameter likely decreased from Mapping 1B to Mapping 2B (Mapping 2B, *M* = 120 ms, 95% *CrI* [94, 145], difference, *M* = 35 ms, 95% *CrI* [5, 66], *pd* = .989). This decrease could be mainly driven by an earlier onset of the goal-directed process in the second assessment (Mapping 1B, *M* = 553 ms, 95% *CrI* [538, 568]; Mapping 2B, *M* = 531 ms, 95% *CrI* [517, 545]; difference, *M* = 22 ms, 95% *CrI* [8, 36], *pd* = .999), whereas the evidence for differences in the speed of the habitual process was uncertain (Mapping 1B, *M* = 398 ms, 95% *CrI* [381, 416]; Mapping 2B, *M* = 411 ms, 95% *CrI* [388, 435]; difference, *M* = -12 ms, 95% *CrI* [-40, 14], *pd* = .821). This supported improved goal-directed performance in the second assessment due to practice effects.

Similar to the habit strength parameter, individual differences in the speed difference parameter were temporally stable in both experiments (Exp. 1, 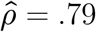, 95% *CrI* [.50, .98]; Exp. 2, 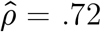, 95% *CrI* [.51, .88]; Fig. 4e). The remaining parameters were similarly stable within each experiment (Supplementary Fig. 6; see Supplementary Fig. 7 for correlations between parameters).

### Training improvement is associated with habitual behavior

We explored whether the extent to which participants automate their actions during training is associated with greater habit expression. We fitted a hierarchical linear model to RTs for correct responses during training, with training day as a predictor, and extracted participant-specific random slopes to measure RT improvement. In Exp. 1, RT improvement across the three days of training was not correlated with habitual errors on the test session (*r*_77_ = -.12; *P* = .278). However, in Exp. 2, which included an additional day of training, RT improvement for each set of associations was associated with habitual errors on the corresponding test (stimulus set 1: *r*_168_ = -.34; *P* < .001; *P*-holm < .001; stimulus set 2: *r*_168_ = -.23; *P* = .001; *P*-holm = .005; Fig. 5a). Notably, RT improvement during neither training phase was associated with other errors (stimulus set 1: *r*_168_ = -.03; *P* = .637; *P*-holm = .637; stimulus set 2: *r*_168_ = -.08; *P* = .269; *P*-holm = .539), indicating that improvements during training were specifically related to habitual responding.

**Figure 5:**
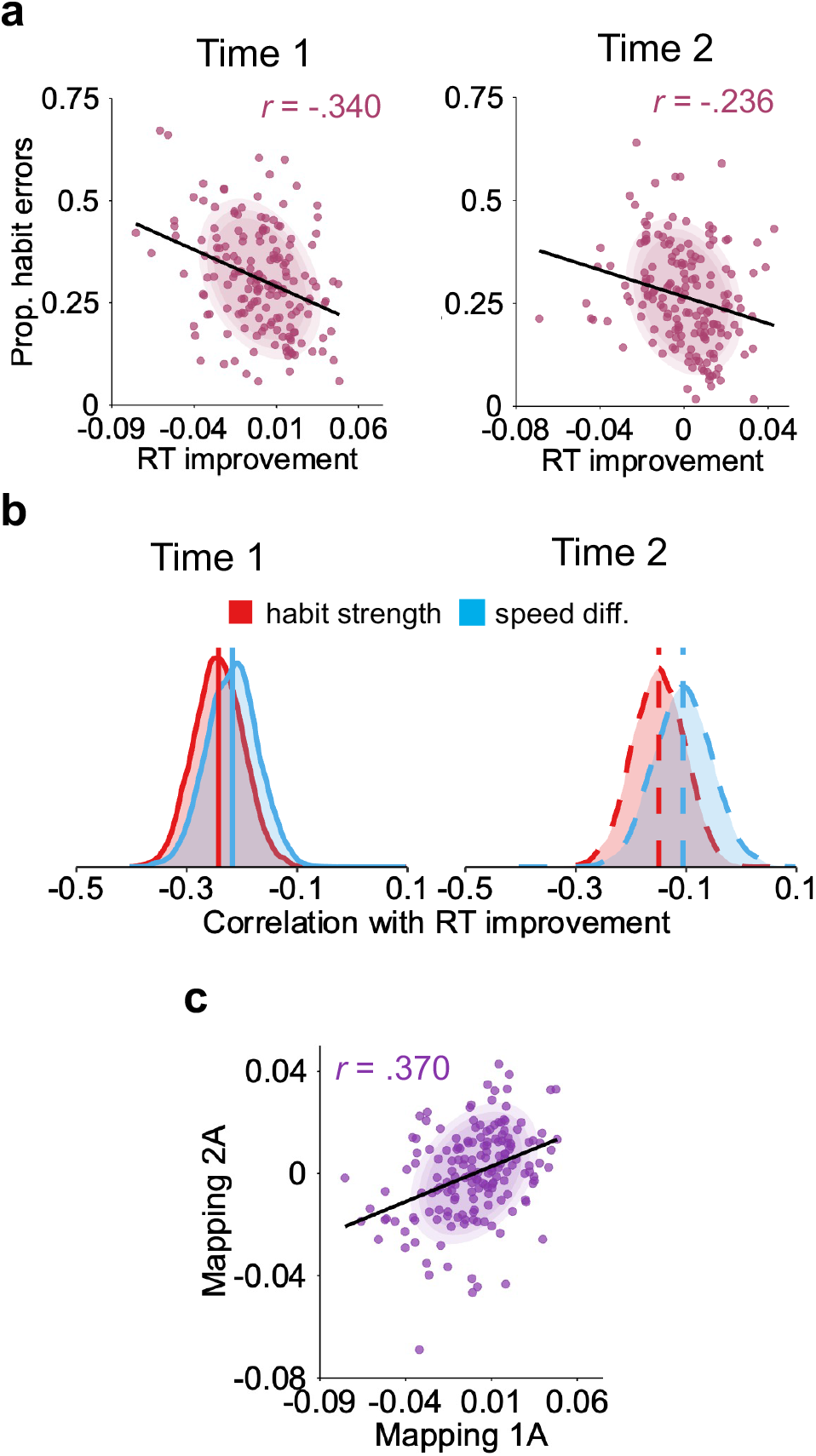
Performance during training was associated with habitual behavior. **(a)** Correlation between improvement in RTs for correct responses during training and the proportion of habitual errors in the corresponding forced-response test. We quantified RT improvement using participant-specific slope coefficients from a hierarchical linear model: *log*(RT) ∼ training day. The left panel shows the association between RT improvement during training on mapping 1A and habitual errors under Mapping 1B (Time 1); the right panel shows the corresponding association for Mappings 2A and 2B (Time 2). **(b)** Posterior distribution of the correlation between improvement in RTs for correct responses during training and the habit strength and speed difference parameters, computed for each posterior draw. **(c)** Correlation between the improvements in RTs for correct responses during training from learning each set of stimulus-response associations (with a 4-month interval between them).

We also explored correlations between RT improvement and the habit strength and speed difference parameters. The posterior correlation of habit strength was negative in both stimulus sets (stimulus set 1: *M* = -.24, 95% *CrI* [-.32, -.15]; stimulus set 2: *M* = -.15, 95% *CrI* [-.24, -.05]; Fig. 5b). Similarly, the relative speed advantage in selecting habitual over goal-directed responses, µ_Δ_, was also associated with RT training improvements in both stimulus sets (stimulus set 1: *M* = -.21, 95% *CrI* [-.31, -.12]; stimulus set 2: *M* = -.10, 95% *CrI* [-.21, -.0001]). Thus, there was evidence that RT improvements during training were associated with both an increased likelihood of selecting a habitual response and a reduction in the overlap between the speed at which the habitual and goal-directed processes operate.

While our findings are consistent with a recent study that also reported a relationship between RT improvement and habit errors^23^, we extend this work by showing that individual differences in action automatization during training remained relatively stable over four months (*r*_168_ = .37; *P* < .001; *P*-holm < .001; Fig. 5c). Taken together, these findings suggest that performance during stimulus–response practice may provide an early signature of a trait-like capacity for habit formation.

## Discussion

The premise that individual differences in how people form and rely on habits contribute to the development and maintenance of several mental disorders has shaped recent theoretical and empirical research in both healthy and clinical populations^17^. However, it remained unclear to what extent these individual differences reflect stable, trait-like tendencies. We provide evidence that human habitual behavior reflects two temporally stable cognitive processes: habit learning, defined as the rate at which a habit is acquired through repetition, and habit expression, defined as the extent to which the habit is performed when it is no longer appropriate. We demonstrated this stability across two preregistered test-retest experiments examining stimulus–response habits. In both experiments, action repetition (Exp. 1: 3 days, 3,000 trials; Exp. 2: 4 days, 4,000 trials) induced habit formation, as evidenced by a greater frequency of habitual errors than other non-practiced errors. Habitual behavior decreased at the second assessments of both experiments. This reduction would, if anything, be expected to attenuate consistency at the individual level. Nevertheless, individual differences in habitual behavior remained stable across assessments, underscoring the robustness of these cognitive processes as trait-like characteristics. Our computational modelling further supported the temporal stability of individual differences in habitual control, showing that both the strength with which habits were expressed and the difference between the preparation of habitual and goal-directed responses were stable.

These findings strengthen theories that conceptualize disorders such as addiction and compulsive disorders, at least in part, as arising from a neurocognitive imbalance between goal-directed and habitual control^9,37,38^. More broadly, our study lends credibility to previous findings linking performance on behavioral habit tasks to diagnosed psychiatric conditions and self-reported psychiatric symptoms^11–13^. These frameworks often raise the question of whether differences in habitual control precede the onset of psychiatric symptomatology, or emerge as their consequence, particularly given that environmental factors such as repeated drug exposure and stress can themselves modify the habit system^9,39^. Our findings help advance investigations into this question. Specifically, experimentally derived measures of habitual behavior can provide stable individual markers, allowing performance to be compared against normative reference values and supporting mechanistic investigations of psychiatric conditions.

The forced-response task provides a habit measure that is sensitive to the amount of training, effectively mapping the acquisition of habits predicted in most animal studies^14,15,18^. Our computational modelling can account for the gradual formation of habits as a function of the amount of training^40–42^, as evidenced by differences in the habit strength parameter between Exp. 1 and Exp. 2. This contrasts with the computational model used by Hardwick et al.^19^. That study accounted for behavioral differences between a group with 4 days of training and another with 20 days of training using the all-or-nothing model. The authors argued that including a habit strength parameter did not offer computational advantages in accounting for their data, and that group differences in habit errors were more parsimoniously explained by differences in the timing at which habitual and goal-directed responses were selected. In contrast, we found evidence that habit strength increased after an additional day of training. There are substantial differences between Hardwick et al’s study and the present work that could explain these divergent computational results. The most notable is the amount of training of the most trained condition (4 vs 20 days). It could be possible that the habit strength and practice do not follow a linear relationship. For instance, it is plausible that habit strength in behavioral tasks rapidly reaches its asymptote (see also ref.^43^). Interestingly, animal research has recently suggested that actions transition abruptly from goal-directed to habitual^44^. Future research in humans may more systematically explore different levels of practice to elucidate how habit commissions and computational parameters evolve in function of training.

Temporal stability in habitual behavior was evident not only in action control after the associations had changed but also during the acquisition of stimulus–response habits. The rate at which participants accelerated their responses during practice remained relatively stable over four months. Thus, although trial-and-error learning varied across individuals, individual learning patterns were consistent over time. Greater improvement during training was linked with greater subsequent action inflexibility during testing. Computationally, training improvement was associated with an increase in habit strength, as well as a larger estimated separation between the habitual and goal-directed onsets. This finding supports the computational rationale for a capacity for habit formation that increases decision-making efficiency, albeit at the cost of greater behavioral inflexibility^45^. In other words, a tendency toward habitual control was detectable even when current goals were aligned with the habitual response. Future research linking these early behavioral markers of habit formation to the development of clinical or subclinical behaviors could determine whether ongoing habitualization predicts their subsequent persistence, potentially informing efforts to prevent maladaptive habits.

Once a habit has been acquired, several strategies have been proposed to disrupt it^17,46^, although no clear consensus has emerged regarding their efficacy. This gap stems partly from the limited experimental evidence directly examining how stimulus–response habits can be weakened. In Exp. 1, we found that habit expression was substantially reduced after practicing competing stimulus–response associations, increasing the probability of responding correctly. Notably, our computational modelling suggests that this decrease in habitual behavior was driven by a reduction in habit strength as well as a faster selection of the goal-directed response during the retest session. This finding indicates that, although the capacity to acquire and control habits may be temporally stable, habitual behavior itself remains malleable. It is important to note, however, that Exp. 1 was designed to study the temporal stability of individual differences in habits, and it cannot clearly disentangle the contribution of the passage of time (∼48 hours since the last training trial) from counterpractice. Nevertheless, this finding may provide preliminary experimental evidence supporting habit substitution interventions. This approach bases behavioral change on training an incompatible response in the same triggering context, as a potentially effective, theory-driven strategy for nonpharmacological intervention^47,48^. The success of habit disruption may depend on identifying and training an alternative stimulus–response association that is genuinely incompatible with the associative structure of the target habit, which may be challenging in real-world settings.

Prior work has shown limited agreement across habit assessments conducted using different instruments and at different time points^20,49^. Although these studies attributed the divergent findings to the possibility that habitual control comprises distinct cognitive processes, they also acknowledged that temporal instability in the construct itself could contribute to the lack of convergence across methods. Our study provides evidence against this instability account, thereby strengthening support for a multidimensional conceptualization of habits. Indeed, action control is increasingly understood as a multi-dimensional process in which different components may habitualize independently (e.g., deciding whether to take an action versus which action to take)^22^, and may be supported by distinct underlying associative structures (including stimulus–response, stimulus–outcome, or stimulus–stimulus)^50–52^. These habitual pathways are also likely to be modulated by both the valence of the behavior (e.g., approach, avoidance, or value-neutral behavior)^53–55^ and the training protocols used^56^. Different behaviors may therefore engage motor and motivational components to varying degrees and differ in how they become habitual. Future research should examine, with greater granularity, which processes involved in habit formation and expression exhibit trait-like stability over time.

Our study provides evidence that individual differences in how humans acquire and express habits reflect stable, trait-like cognitive processes. These findings not only strengthen neurocognitive accounts implicating habitual processes in psychopathology but also provide a foundation for investigating how genetic, neural, and environmental factors shape these processes. More broadly, these findings open new avenues for developing and refining measures of habit propensity and may ultimately inform novel approaches to the diagnosis, prevention, and treatment of psychiatric disorders.

## Methods

### Preregistrations

The experiments reported in the present study were preregistered before data collection (preregistration documents: Exp. 1, https://doi.org/10.17605/OSF.IO/SQ5GU (ref.^57^); Exp. 2, https://doi.org/10.17605/OSF.IO/B4A9J (ref.^58^)). For each experiment, the preregistered hypotheses, design, sampling plan, variables and planned analyses are publicly available in the corresponding preregistration document.

### Planned sample sizes

Power analyses were conducted assuming 95% statistical power and an alpha level of .05 for one-tailed Pearson’s *r* tests. In Exp. 1, we aimed to detect a latent effect size of *r* = .50. To determine the target effect size, we accounted for the impact of measurement error on statistical power and estimated the expected attenuation of that effect size^27,59^. Specifically, we calculated the split-half permutation-based reliability of the primary measures using the publicly available dataset from Exp. 1 of Hardwick et al.^19^, which used a similar amount of training and a comparable task version. The reliability estimate of our main measure in that dataset was .75. Using Spearman’s attenuation formula^28^, we reduced the target effect size accordingly. This yielded an observed target effect size of *r* = .375 for the power analysis, which indicated a required sample size of 72 participants. In Exp. 2, the sample size was determined assuming an effect size 50% smaller than that observed in Exp. 1 (*r* = .59 × .50 = .295), because the test–retest assessments involved different stimuli and were separated by a 4-month interval. The power analysis indicated that 117 participants were required. To account for potential attrition and exclusions, 126 and 260 participants were invited to participate in Exp. 1 and Exp. 2, respectively.

### Participants

A total of 313 participants took part in this study. All were at least 18 years old, were Spanish speakers, had normal or corrected-to-normal vision, and reported no known neurological disorders. All participants provided written informed consent. Participants in Exp. 1 received €5 and course credit, whereas those in Exp. 2 received course credit. Exp. 1 included 91 participants (mean age = 20.65 years, standard deviation (*SD*) = 3.30; 73 women), and Exp. 2 included 222 participants (mean age = 20.48 years, *SD* = 5.34; 189 women). Participants were recruited through the University of Málaga and, for Exp. 2, additionally through the Autonomous University of Madrid website. The experimental procedures were approved by the ethics committees of the University of Málaga (84-2025-H) and the Autonomous University of Madrid (CEI-144-3228).

### Exclusion criteria

Preregistered exclusion criteria were applied in both experiments. One participant in Exp. 1 and 19 participants in Exp. 2 did not complete all experimental sessions (five and 10 sessions, respectively) and were therefore excluded. One additional participant in Exp. 1 and two in Exp. 2 were excluded due to a technical error during one of the sessions. Preregistered trial-level exclusions were applied to the forced-response blocks. We excluded trials in which participants responded before stimulus onset (anticipations: Exp. 1: 2.68 %; Exp. 2: 3.01%) or failed to respond (timeouts: Exp. 1: 1.11%; Exp. 2: 1.01%). We also preregistered participant level exclusion criteria, based on Du and Haith^22^, to identify participants who did not follow the task instructions. These criteria focused on performance in the criterion and forced-response blocks. In Exp. 2, one participant was excluded for requiring more than 600 trials to complete a criterion block. For the forced-response blocks, participants were excluded if they had more than 20% anticipations or timeouts (Exp. 1, four participants; Exp. 2, two participants), fewer than 8% of trials with preparation times between 0 and 200 ms, allowing for up to 20% anticipations or timeouts (Exp. 1, four participants; Exp. 2, 12 participants), or accuracy below .50 for any association type at preparation times greater than 900 ms (Exp. 1, two participants; Exp. 2, 16 participants).

### Experimental task

Participants completed blocks of the forced-response task. The task was programmed using Psychtoolbox (v3.0.19.4)^60^ and run in MATLAB 2022b^61^.

Participants placed the index, middle, ring, and little fingers of their dominant hand on the “H,” “U,” “I,” and “L” keys, respectively, with each corresponding to one of four possible symbols. On each trial, a symbol appeared on the screen, and participants were required to press the corresponding key. Stimuli were presented in a pseudo-random order such that each symbol appeared five times within each 20-trial epoch and no symbol appeared more than twice consecutively. Each block comprised 100 trials. Two sets of four stimuli were used, and their assignment was counterbalanced across participants.

During the experiment, two stimulus-key mappings were used: Mapping A and Mapping B. For Mapping A, four stimulus-key associations were assigned randomly to each participant according to a counterbalanced design. Mapping B was a modified version of Mapping A in which the responses associated with two stimuli were switched, resulting in two consistent S-R associations and two remapped S-R associations. Mapping A was always used during the training sessions, whereas Mapping B was used during the test sessions. The response switch in Mapping B occurred either between the “H” and “I” keys or between the “U” and “L” keys, with the two alternatives counterbalanced across participants. Participants completed three types of blocks: criterion, training, and forced-response test blocks.

#### Criterion blocks

Participants learned the four symbol-key associations of Mapping A or Mapping B through trial and error. They were instructed to prioritize accuracy. On each trial, a symbol was displayed at the center of the screen. A correct response turned an on-screen feedback box green and played a pleasant sound. The next trial began after 300 ms. An incorrect response turned the on-screen feedback box red, triggered a buzzer, and prevented further responses for 1 second. The trial ended only after the correct response was made. The block ended once the participant had responded correctly to each stimulus five consecutive times.

#### Training blocks

During training, participants practiced the associations from Mapping A. They were instructed to respond as quickly and accurately as possible. The trial structure was the same as in the criterion blocks. At the end of each block, participants were informed of their completion time and their fastest time in previous blocks. They were encouraged to improve on their best performance.

#### Forced-response test blocks

We used forced-response blocks to characterize the speed–accuracy trade-off as a function of preparation time (i.e., response time minus stimulus onset) for each response type and association. This allowed us to examine time-dependent patterns of habit expression. Participants heard a sequence of four tones separated by 400 ms, with the stimulus appearing at a randomly selected time during the sequence (uniformly distributed across trials). They were instructed to respond according to Mapping B and to time their response to coincide with the fourth tone, which occurred 1,200 ms after trial onset.

Correct and incorrect responses changed the corresponding on-screen feedback box to the appropriate color. The next trial began after 300 ms. Responses occurring between 100 ms before and 75 ms after the fourth tone elicited a “good timing” message at the center of the screen. Earlier responses elicited the message “a bit early,” whereas later responses elicited the message “a bit late.” Responses occurring more than 300 ms before or more than 200 ms after the fourth tone elicited the message “too early” or “too late,” respectively. Participants then incurred a 2-s penalty and had to press any key to proceed to the next trial. If no response was made within 1.5 s, the message “too late” was displayed and the same penalty was applied.

### Procedure

In Exp. 1, participants attended the laboratory in person on five consecutive days in April 2025. The first session began with a 100-trial familiarization training block using arbitrary stimuli depicting colored fingers, followed by a criterion block with Mapping A and 10 training blocks. On Days 2 and 3, participants completed 10 training blocks per day. On Days 4 and 5, they completed forced-response familiarization blocks, followed by a criterion block (Mapping B), and five forced-response test blocks. Participants completed two to four forced-response familiarization blocks, depending on their performance. If more than 20% of responses occurred either more than 300 ms before or more than 200 ms after the fourth tone, participants completed an additional familiarization block, up to a maximum of four blocks. Each block comprised 100 trials.

In Exp. 2, participants completed five consecutive in-person laboratory sessions in November 2025 and repeated the same 5 day sequence with a different stimulus set in March 2026 (ten sessions in total). The first session of each 5 day sequence began with a familiarization training block, followed by a criterion block (Mapping A) and 10 training blocks. Each of the next three sessions comprised 10 training blocks. In the fifth session, participants completed two to four forced-response familiarization blocks, following the same performance-based procedure as in Exp. 1, followed by a criterion block with Mapping B and five forced-response test blocks.

### Data analysis

#### Criterion blocks

The number of trials required to reach the criterion terminating each block was recorded as a measure of the ability to acquire the relevant set of stimulus-response associations (Mapping A or Mapping B).

#### Training blocks

Response speed was indexed by reaction time (RT), defined as the interval between trial onset and the participant’s response. RT analyses included only trials on which the first response was correct. Accuracy was indexed by the error rate, calculated as the number of errors per block.

#### Forced-response blocks

For each response type and association type (consistent or remapped), we constructed a speed–accuracy trade-off as a function of preparation time. For both association types, we recorded RTs and correct responses. For remapped associations, we additionally classified habitual responses as responses that had been correct under Mapping A but became incorrect under Mapping B following the response switch. Other errors were defined as incorrect responses that did not correspond to the habitual response. Preparation time was calculated as RT minus stimulus onset time. Time-dependent response patterns were then examined using a 100-ms sliding window, advanced in 1-ms increments, within which the proportion of each response type was calculated separately for consistent and remapped associations.

### Statistical analysis

Data processing, statistical analyses, and visualization were conducted using RStudio (v. 2024.12.1)^62^.

#### Group-level effects

Group-level effects were analyzed using two-tailed *t*-tests, except for preregistered one-tailed *t*-tests, as specified in the main text. When comparing two independent samples, we used Welch’s two-sample *t*-test, which does not assume equal variances.

#### Temporal stability of individual differences in habitual errors

The temporal stability of individual differences in habitual responding was assessed using preregistered one-tailed Pearson’s r correlations between the proportion of habitual errors between test sessions. To estimate the psychometric reliability of the measure in each session, we calculated split-half permutation-based reliability using the *rapidsplithalf* R package^63^. Across 10,000 iterations, trials were randomly assigned to two halves, and Pearson’s *r* was calculated between the halves and adjusted using the Spearman–Brown prediction formula^64^. The mean adjusted coefficient, *r*_sb_, served as the final reliability estimate, with the 2.5th and 97.5th percentiles defining the 95% bootstrap confidence interval.

To facilitate interpretation of the temporal stability results, we accounted for measurement error by fitting a generative model to habitual responding data from both sessions^29^. A Bayesian hierarchical model was implemented using the *brms* package in R^65^, with *cmd-stanr* as the sampling backend. Habit errors were modeled using a Bernoulli distribution with a logit link. Session was entered as a fixed effect, with the population-level intercept omitted. The random-effects structure likewise omitted the intercept, allowing separate participant-level random coefficients to be estimated for each session. The model formula was as follows:

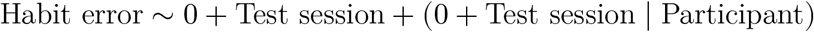

Under this parameterization, the correlation between the participant-level coefficients for the two sessions provides a latent, deattenuated estimate of temporal stability. In the main text, we report the posterior mode of this correlation together with its 95% credible interval (*CrI*). This procedure closely approximates Spearman’s attenuation correction formula^28^, while providing an estimate of uncertainty around this value^29,30^.

For each experiment, the model was fitted using four chains, each with 500 warm-up iterations and 4,500 post warm-up sampling iterations. Weakly informative priors were specified. Fixed-effect coefficients were assigned normal priors centered at zero with an SD of 1 on the logit scale. The standard deviations of the random effects were assigned corresponding half-normal priors, and the correlation structure of the random effects was modeled using an LKJ prior with shape parameter *η* = 1, which is uniform over the space of valid correlation matrices^66^. For visualization, participant-level posterior means were transformed from the logit scale to the probability scale.

#### Model fitting procedure

We fitted a no-habit, a continuum and an all-or-nothing habit model following the computational framework described by Hardwick et al.^19^ (see *Models of habit formation*). We used the R package *cmdstanr* (v. 0.9.0)^67^ to fit the models in Stan^68^. Posterior distributions were estimated using Hamiltonian Monte Carlo with the No-U-Turn Sampler (NUTS), running four chains with 1,000 warm-up iterations and 3,000 post-warm-up sampling iterations per chain. The adapt_delta parameter was set at .95. We required 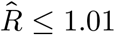, bulk- and tail-effective sample size ≥ 400 for all parameters, and the Bayesian fraction of missing information ≥ 0.2 for all chains, with no divergent transitions observed^69^.

When necessary, the posterior distributions of each parameter were transformed to their original scale (i.e., back-log-transformation or inverse-logit-transformation) (see below). From those distributions, we report the mean value, the 95% credible interval (*CrI*), and the probability of direction (*pd*; range: 0.5 to 1.0). The *pd* refers to the proportion of posterior samples sharing the sign of the posterior median; a *pd* higher than .95 approximately corresponds to a one-sided *P*-value lower than .05. We interpreted *pd* values following common recommendations and terminology (e.g., likely, uncertain)^70,71^.

#### Model comparison

To determine which model best fit our data, we compared the models based on their out-of-sample predictive performance using expected log pointwise predictive density (ELPD) estimates obtained via Pareto-smoothed importance sampling leave-one-out cross-validation (PSIS-LOO)^33^. If any Pareto *k* values exceeded 0.7, we applied moment matching^72^. We concluded that a model accounted for our data better if its ELPD difference with the second-best model exceeded twice the standard error of that difference^33^.

#### Association between training improvement and habitual behavior

We defined improvement during training as the reduction in reaction times (RTs) for correct responses across training sessions. For each training phase, we filtered for trials in which the first response was correct and fitted a hierarchical linear model to the log-transformed RTs to better approximate normality. Training session was included as both a fixed effect and a participant-level random slope. The model was fitted using the *lme4* R package^73^ and specified as follows:

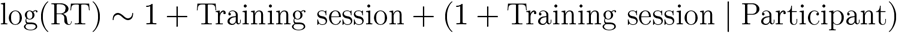

We extracted participant-specific slopes as an index of RT improvement during training. These slopes were then correlated (two-sided Pearson’s *r*) with habitual errors, other errors, and RT improvement across training phases. To control the family-wise error rate at *α* = 0.05, we applied the Holm correction. Additionally, we correlated RT improvements with the habit strength and speed difference parameters estimated in their corresponding forced-response blocks (see below). The correlation was computed for each posterior draw, hence generating a posterior distribution of the correlation.

#### Models of habit formation

We followed the computational framework developed by Hardwick et al.^19^ for the forced-response task to formalize three theoretical models of habit formation: the no-habit model, the all-or-nothing model and the continuum model. This framework estimates the speed at which habitual (i.e., trained responses for revised stimuli) or goal-directed responses (i.e., correct responses for revised stimuli) are prepared at independent random times, 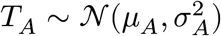 and 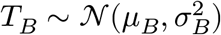, respectively. It is assumed that the goal-directed Gaussian distribution has a greater mean than that of the habitual distribution (µ_*A*_ < µ_*B*_). If no response was available by the time of responding, *t* (i.e., *t* < *T*_*A*_), around chance performance would be expected; if only the habitual response has been prepared (i.e., *t* > *T*_*A*_), it may be expressed; once the goal-directed response has been prepared (i.e., *t* > *T*_*B*_), the correct response would be expressed.

Following the matrix formulation of Du and Haith^22^, the models can be mathematically expressed in a compact form as a function of three matrices:

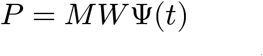

where Ψ(*t*) attributes the probability of each event (selecting a habitual response or a goal-directed one) as a time-dependent vector, while *M* captures the probability of each response type conditional on which event has occurred or not. *W* introduces moderators for habitual and goal-directed responses.

The vector Ψ(*t*) is as follows:

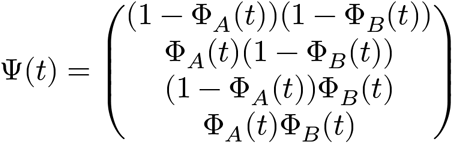

From top to bottom, the matrix rows represent the possible events at time *t*: none of the responses was prepared, only the original response was prepared, only the revised response was prepared, and both responses were prepared. 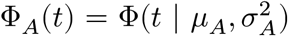 and 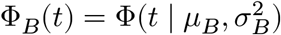 represent the cumulative distribution of *T*_*A*_ and *T*_*B*_, respectively.

The matrix *M* is as follows:

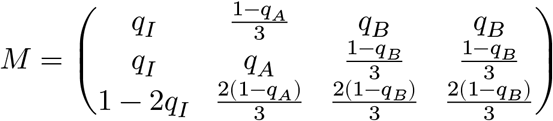

The rows of the matrix represent the probability of emitting each response type, from top to bottom: goal-directed (*q*_*B*_; correct), habitual (*q*_*A*_; original) or another response never associated with the stimuli (*q*_*I*_; other), conditional on the occurrence of each event. Each column corresponds to one of the events listed in the previous matrix, taken in top-to-bottom order from left to right.

The matrix *W* is as follows:

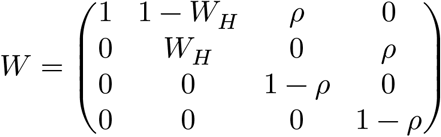

Matrix *W* introduces parameters that modulate the probability of event occurrence (none of the responses prepared, habitual response only, goal-directed response only, both responses). *W*_*H*_ is habit strength and modulates the probability that a habit error would be emitted if it had been prepared. Similarly, *ρ* is the lapse rate of the goal-directed response, representing the probability that it is not emitted even when available.

The continuum model included all the parameters described above, Θ = (µ_*A*_, σ_*A*_, µ_*B*_, σ_*B*_, *q*_*A*_, *q*_*B*_, *q*_*I*_, *W*_*H*_, *ρ*). As in previous studies^19,22^, *q*_*A*_ was fixed at .95, reflecting the typical success rate for well-learned associations, because it is difficult to estimate in practice. Thus, eight parameters were estimated. In addition, we reparametrized µ_*B*_ as µ_*B*_ = µ_*A*_ + µ_Δ_ with µ_Δ_ > 0, which ensures that µ_*A*_ < µ_*B*_ but also improves sampling^74^. In the all-or-nothing model, when available, the habitual response is always expressed unless overridden by the goal-directed response. This is equivalent to setting *W*_*H*_ = 1, so seven parameters were estimated. In the no-habit model, we assumed that habits were not learned, so only the revised association can be available. We implemented this by setting *W*_*H*_ = 0 and *ρ* = 0, and by making the two columns of *M* that differ only by the habitual response identical. This model therefore has only four free parameters, Θ = (µ_*B*_, σ_*B*_, *q*_*B*_, *q*_*I*_).

Parameters were defined hierarchically across participants and sessions. For each parameter, participant-level values in the two sessions followed a bivariate normal distribution on the unconstrained scale with session-specific group means and standard deviations and a between-session correlation, which was our estimate of temporal stability. While we supported our prior specification with participant-level estimates reported in previous research (i.e., in their parameter recovery analyses^19,22^), we used regularizing priors that allow a broad range of parameter values but disallow implausible ones. We used the same priors for both sessions (Table 2; see Supplementary Fig. 3 for prior predictive checks). We used a non-centered parameterization and LKJ priors with shape parameter *η* = 1 to ensure uniform expectation across all correlation values, and we Cholesky-factorized them to aid estimation.

**Table 2:** Priors for each parameter. Normal^+^ denotes a half-normal distribution. Except for the no-habit model, µ_*B*_ was reparametrized as µ_*A*_ + µΔ. Correct and habit errors were mapped to one key and other errors to two; hence, *q*_*I*_ was scaled by 0.5.

| Parameter | Estimated on | Location | Scale |
|---|---|---|---|
| *μ*<sub>*A*</sub> | identity (s) | Normal(0.45, 0.1) | Normal<sup>+</sup>(0, 0.1) |
| *μ*<sub>Δ</sub> | log | Normal(-2.3, 0.3) | Normal<sup>+</sup>(0, 0.5) |
| *μ*<sub>*B*</sub> (no-habit model only) | identity (s) | Normal(0.55, 0.1) | Normal<sup>+</sup>(0, 0.1) |
| *σ*<sub>*A*</sub> | log | Normal(-2.3, 0.2) | Normal<sup>+</sup>(0, 0.3) |
| *σ*<sub>*B*</sub> | log | Normal(-2.3, 0.2) | Normal<sup>+</sup>(0, 0.3) |
| *q*<sub>*B*</sub> | logit | Normal(1.5, 0.5) | Normal<sup>+</sup>(0, 1) |
| *q*<sub>*I*</sub> | logit | Normal(0, 1) | Normal<sup>+</sup>(0, 1) |
| *W*<sub>*H*</sub> | logit | Normal(0, 1) | Normal<sup>+</sup>(0, 0.6) |
| *ρ* | logit | Normal(-2.5, 0.5) | Normal<sup>+</sup>(0, 0.6) |
| Correlations | – | LKJ(1) | – |

## Data availability

All data and materials are available via Zenodo at https://doi.org/10.5281/zenodo.23926926 (ref.^75^).

## Code availability

All code is available via Zenodo at https://doi.org/10.5281/zenodo.23926926 (ref.^75^).

## Author contributions

Conceptualization: PML and DL. Methodology: PML, DL, FGF, AVM, JJL, SM, MAV and TGF. Software: PML. Investigation: PML, SM, AVM, TGF, MAV and JJL. Data curation: PML. Formal analysis: PML, JJL and FGF. Validation: PML. Visualization: PML and FGF. Funding acquisition: DL. Project administration: DL. Supervision: DL. Writing – original draft: PML. Writing – review & editing: DL, JJL, FGF, AVM, SM, TGF and MAV.

## Supplementary information

Supplementary information is available via Zenodo at https://doi.org/10.5281/zenodo.23926747 (ref.^76^).

## References

1. Lally, P., Van Jaarsveld, C. H. M., Potts, H. W. W. & Wardle, J. How are habits formed: Modelling habit formation in the real world. European Journal of Social Psychology 40, 998–1009 (2010).

2. Lin, P.-Y., Wood, W. & Monterosso, J. Healthy eating habits protect against temptations. Appetite 103, 432–440 (2016).

3. Bayer, J. B., Anderson, I. A. & Tokunaga, R. S. Building and breaking social media habits. Current Opinion in Psychology 45, 101303 (2022).

4. Turner, G. et al. A computational model of reward learning and habits on social media. Nature Communications 17, 7170 (2026).

5. Dickinson, A., Balleine, B., Watt, A., Gonzalez, F. & Boakes, R. A. Motivational control after extended instrumental training. Animal Learning & Behavior 23, 197–206 (1995).

6. Wit, S. de & Dickinson, A. Associative theories of goal-directed behaviour: A case for animal–human translational models. Psychological Research 73, 463–476 (2009).

7. Crum, K. L. et al. Leveraging Habits to Improve Adherence to Gout Medications: A Qualitative Study. ACR Open Rheumatology 6, 625–633 (2024).

8. Verplanken, B., Aarts, H. & Van Knippenberg, A. Habit, information acquisition, and the process of making travel mode choices. European Journal of Social Psychology 27, 539–560 (1997).

9. Everitt, B. J. & Robbins, T. W. Drug Addiction: Updating Actions to Habits to Compulsions Ten Years On. Annual Review of Psychology 67, 23–50 (2016).

10. Gillan, C. M. Recent Developments in the Habit Hypothesis of OCD and Compulsive Disorders. in The Neurobiology and Treatment of OCD: Accelerating Progress vol. 49 (Springer International Publishing, Cham, 2021).

11. Sookud, S., Martin, I., Gillan, C. M. & Wise, T. Impaired Goal-Directed Planning in Transdiagnostic Compulsivity Is Explained by Uncertainty About Learned Task Structure. Biological Psychiatry: Cognitive Neuroscience and Neuroimaging 11, 357–365 (2026).

12. Banca, P. et al. Action-sequence learning, habits and automaticity in obsessive-compulsive disorder. eLife 12, RP87346 (2024).

13. Donegan, K. R. et al. Compulsivity is associated with accelerated stimulus-response habit learning. (2025) doi:10.31234/osf.io/xwtsu_v1.

14. Martínez-López, P., Garre-Frutos, F., Vadillo, M. A. & Luque, D. The Experimental Study of Habitual Behavior in Humans: A Review of the Validity of Current Protocols. (2026) doi:10.31234/osf.io/r62pa_v1.

15. Balleine, B. W. & O’Doherty, J. P. Human and Rodent Homologies in Action Control: Corticostriatal Determinants of Goal-Directed and Habitual Action. Neuropsychopharmacology 35, 48–69 (2010).

16. Dolan, R. J. & Dayan, P. Goals and Habits in the Brain. Neuron 80, 312–325 (2013).

17. Buabang, E. K., Donegan, K. R., Rafei, P. & Gillan, C. M. Leveraging cognitive neuroscience for making and breaking real-world habits. Trends in Cognitive Sciences 29, 41–59 (2025).

18. Watson, P., O’Callaghan, C., Perkes, I., Bradfield, L. & Turner, K. Making habits measurable beyond what they are not: A focus on associative dual-process models. Neuroscience & Biobehavioral Reviews 142, 104869 (2022).

19. Hardwick, R. M., Forrence, A. D., Krakauer, J. W. & Haith, A. M. Time-dependent competition between goal-directed and habitual response preparation. Nature Human Behaviour 3, 1252–1262 (2019).

20. Martínez-López, P., Vázquez-Millán, A., Garre-Frutos, F. & Luque, D. Assessing the validity evidence for habit measures based on time pressure. Behavior Research Methods 58, 7 (2026).

21. Rafei, P., Suddell, S. F., Buabang, E. K., Urcelay, G. P. & Gillan, C. M. Does time pressure reveal habits? Divergent results from outcome devaluation and response remapping paradigms. Cognitive, Affective, & Behavioral Neuroscience https://doi.org/10.3758/s13415-026-01474-6 (2026) doi:10.3758/s13415-026-01474-6.

22. Du, Y. & Haith, A. M. Dissociable habits of response preparation versus response initiation. Nature Human Behaviour 9, 1941–1958 (2025).

23. Buabang, E. K. et al. Neural dynamics of covert habit activation and control in humans. (2025) doi:10.1101/2025.11.12.687990.

24. Kuchibhotla, K. V. et al. Dissociating task acquisition from expression during learning reveals latent knowledge. Nature Communications 10, 2151 (2019).

25. Kelly, A. M. C. & Garavan, H. Human Functional Neuroimaging of Brain Changes Associated with Practice. Cerebral Cortex 15, 1089–1102 (2005).

26. Karvelis, P., Paulus, M. P. & Diaconescu, A. O. Individual differences in computational psychiatry: A review of current challenges. Neuroscience & Biobehavioral Reviews 148, 105137 (2023).

27. Hedge, C., Powell, G. & Sumner, P. The reliability paradox: Why robust cognitive tasks do not produce reliable individual differences. Behavior Research Methods 50, 1166–1186 (2018).

28. Spearman, C. The Proof and Measurement of Association between Two Things. The American Journal of Psychology 15, 72 (1904).

29. Haines, N. et al. A tutorial on using generative models to advance psychological science: Lessons from the reliability paradox. Psychological Methods https://doi.org/10.1037/met0000674 (2025) doi:10.1037/met0000674.

30. Rouder, J. N., Kumar, A. & Haaf, J. M. Why many studies of individual differences with inhibition tasks may not localize correlations. Psychonomic Bulletin & Review 30, 2049–2066 (2023).

31. Schurr, R., Reznik, D., Hillman, H., Bhui, R. & Gershman, S. J. Dynamic computational phenotyping of human cognition. Nature Human Behaviour 8, 917–931 (2024).

32. Adkins, T. J., Zhang, H. & Lee, T. G. People are more error-prone after committing an error. Nature Communications 15, 6422 (2024).

33. Vehtari, A., Gelman, A. & Gabry, J. Practical Bayesian model evaluation using leave-one-out cross-validation and WAIC. Statistics and Computing 27, 1413–1432 (2017).

34. Brown, V. M., Chen, J., Gillan, C. M. & Price, R. B. Improving the Reliability of Computational Analyses: Model-Based Planning and Its Relationship With Compulsivity. Biological Psychiatry: Cognitive Neuroscience and Neuroimaging 5, 601–609 (2020).

35. Haines, N., Sullivan-Toole, H. & Olino, T. From Classical Methods to Generative Models: Tackling the Unreliability of Neuroscientific Measures in Mental Health Research. Biological Psychiatry: Cognitive Neuroscience and Neuroimaging 8, 822–831 (2023).

36. Rey-Sáez, R., Franco-Martínez, A., Revuelta, J. & Vadillo, M. A. A Unified Framework for Psychometrics in Experimental Psychology: The Standardized Generalized Hierarchical Factor Model. Psychological Methods https://doi.org/10.1037/met0000846 (2026) doi:10.1037/met0000846.

37. Robbins, T. W., Gillan, C. M., Smith, D. G., De Wit, S. & Ersche, K. D. Neurocognitive endophenotypes of impulsivity and compulsivity: Towards dimensional psychiatry. Trends in Cognitive Sciences 16, 81–91 (2012).

38. Gillan, C. M. et al. Disruption in the Balance Between Goal-Directed Behavior and Habit Learning in Obsessive-Compulsive Disorder. American Journal of Psychiatry 168, 718–726 (2011).

39. Giovanniello, J. R. et al. A dual-pathway architecture for stress to disrupt agency and promote habit. Nature 640, 722–731 (2025).

40. Luque, D., Molinero, S., Watson, P., López, F. J. & Le Pelley, M. E. Measuring Habit Formation Through Goal-Directed Response Switching. Journal of Experimental Psychology: General 149, 1449–1459 (2020).

41. Grundmann, C. C. et al. Studying human habit formation through motor sequence learning. Cognitive, Affective, & Behavioral Neuroscience 25, 941–951 (2025).

42. Vázquez-Millán, A., Martínez-López, P., Rueda, M., León, J. J. & Luque, D. The evaluation of devaluation: Deficient outcome devaluation leads to wrongly considering goal-directed actions as habits. Behavior Research Methods 58, 224 (2026).

43. Yang, C. S., Cowan, N. J. & Haith, A. M. Control becomes habitual early on when learning a novel motor skill. Journal of Neurophysiology 128, 1278–1291 (2022).

44. Moore, S. et al. Revealing abrupt transitions from goal-directed to habitual behavior. Nature Communications 17, 4751 (2026).

45. Marr, D. Vision: A Computational Investigation into the Human Representation and Processing of Visual Information. (MIT Press, Cambridge, Mass, 1982).

46. Wood, W. & Rünger, D. Psychology of Habit. Annual Review of Psychology 67, 289–314 (2016).

47. Verplanken, B. & Wood, W. Interventions to Break and Create Consumer Habits. Journal of Public Policy & Marketing 25, 90–103 (2006).

48. Gardner, B. et al. Breaking habits or breaking habitual behaviours? Old habits as a neglected factor in weight loss maintenance. Appetite 162, 105183 (2021).

49. Nebe, S., Kretzschmar, A., Brandt, M. C. & Tobler, P. N. Characterizing Human Habits in the Lab. Collabra: Psychology 10, 92949 (2024).

50. Dezfouli, A. & Balleine, B. W. Actions, Action Sequences and Habits: Evidence That Goal-Directed and Habitual Action Control Are Hierarchically Organized. PLoS Computational Biology 9, e1003364 (2013).

51. Buabang, E. K. et al. A Goal-Directed Account of Action Slips: The Reliance on Old Contingencies. Journal of Experimental Psychology: General 152, 496–508 (2023).

52. González, A. & De Brugada, I. Habits in Pavlovian learning: Resistance to devaluation by sensory-specific satiety following extensive exposure to palatable tastes. Psychonomic Bulletin & Review 33, 65 (2026).

53. Guitart-Masip, M., Duzel, E., Dolan, R. & Dayan, P. Action versus valence in decision making. Trends in Cognitive Sciences 18, 194–202 (2014).

54. Kahneman, D. & Amos, T. Prospect Theory: An Analysis of Decision under Risk. Econometrica 47, 263–291 (1979).

55. Miller, K. J., Shenhav, A. & Ludvig, E. A. Habits without values. Psychological Review 126, 292–311 (2019).

56. Smolen, P., Zhang, Y. & Byrne, J. H. The right time to learn: Mechanisms and optimization of spaced learning. Nature Reviews Neuroscience 17, 77–88 (2016).

57. Martínez-Lopez, P. et al. Lingering habits: Testing the temporal stability of habits. (2025) doi:10.17605/OSF.IO/SQ5GU.

58. Martínez-Lopez, P. et al. On the temporal stability of the individual differences in learning and expressing habits. (2026) doi:10.17605/OSF.IO/B4A9J.

59. Parsons, S.Kruijt, A.-W. & Fox, E. Psychological Science Needs a Standard Practice of Reporting the Reliability of Cognitive-Behavioral Measurements. Advances in Methods and Practices in Psychological Science 2, 378–395 (2019).

60. Brainard, D. H. The Psychophysics Toolbox. Spatial Vision 10, 433–436 (1997).

61. Inc., T. M. MATLAB. (2022).

62. Team, R. C. R: A Language and Environment for Statistical Computing. (2023).

63. Kahveci, S. Rapidsplithalf: A Fast Permutation-Based Split-Half Reliability Algorithm. (2024) doi:10.32614/CRAN.package.rapidsplithalf.

64. Spearman, C. Correlation calculated from faulty data. British Journal of Psychology 3, 271–295 (1910).

65. Bürkner, P.-C. Advanced Bayesian Multilevel Modeling with the R Package brms. The R Journal 10, 395 (2018).

66. Lewandowski, D., Kurowicka, D. & Joe, H. Generating random correlation matrices based on vines and extended onion method. Journal of Multivariate Analysis 100, 1989–2001 (2009).

67. Gabry, J., Češnovar, R., Johnson, A. & Bronder, S. Cmdstanr: R Interface to ‘CmdStan’. (2025).

68. Carpenter, B. et al. Stan: A probabilistic programming language. Journal of statistical software 76, (2017).

69. Baribault, B. & Collins, A. G. E. Troubleshooting Bayesian cognitive models. Psychological Methods 30, 128–154 (2025).

70. Makowski, D., Ben-Shachar, M. S., Chen, S. H. A. & Lüdecke, D. Indices of Effect Existence and Significance in the Bayesian Framework. Frontiers in Psychology 10, 2767 (2019).

71. Ben-Shachar, M. S., Lüdecke, D. & Makowski, D. Effectsize: Estimation of Effect Size Indices and Standardized Parameters. Journal of Open Source Software 5, 2815 (2020).

72. Paananen, T., Piironen, J., Bürkner, P.-C. & Vehtari, A. Implicitly adaptive importance sampling. Statistics and Computing 31, 16 (2021).

73. Bates, D., Mächler, M., Bolker, B. & Walker, S. Fitting Linear Mixed-Effects Models Using lme4. Journal of Statistical Software 67, (2015).

74. Gelman, A., Van Dyk, D. A., Huang, Z. & Boscardin, J. W. Using Redundant Parameterizations to Fit Hierarchical Models. Journal of Computational and Graphical Statistics 17, 95–122 (2008).

75. Martínez-López, P. et al. Data, materials and code for “temporal stability for individual differences in both habit learning and expression”. (2026) doi:10.5281/zenodo.23926926.

76. Martínez-López, P. et al. Supplementary information for “temporal stability for individual differences in both habit learning and expression”. (2026) doi:10.5281/zenodo.23926747.

