## Supplementary information for "Temporal stability for individual differences in both habit learning and expression"

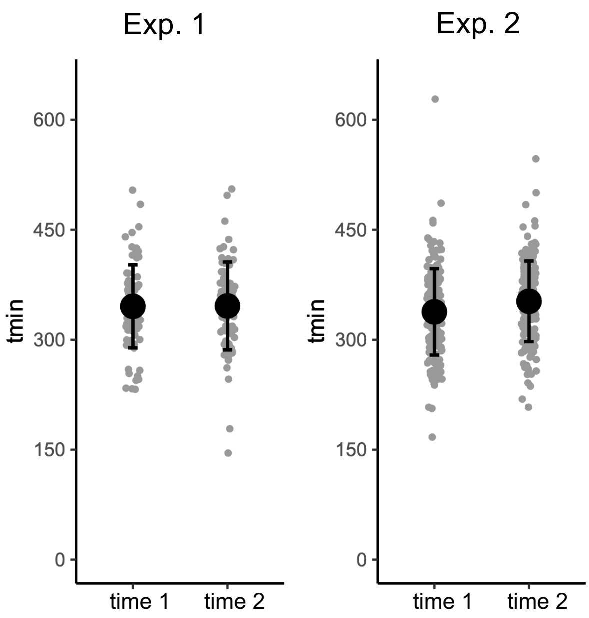


**Supplementary Figure 1: Estimates of tmin in each experiment and test session.** Grey dots indicate participant-level values. Black dots represent session means in each experiment, and error bars show standard deviations (SD). Values are presented in milliseconds.

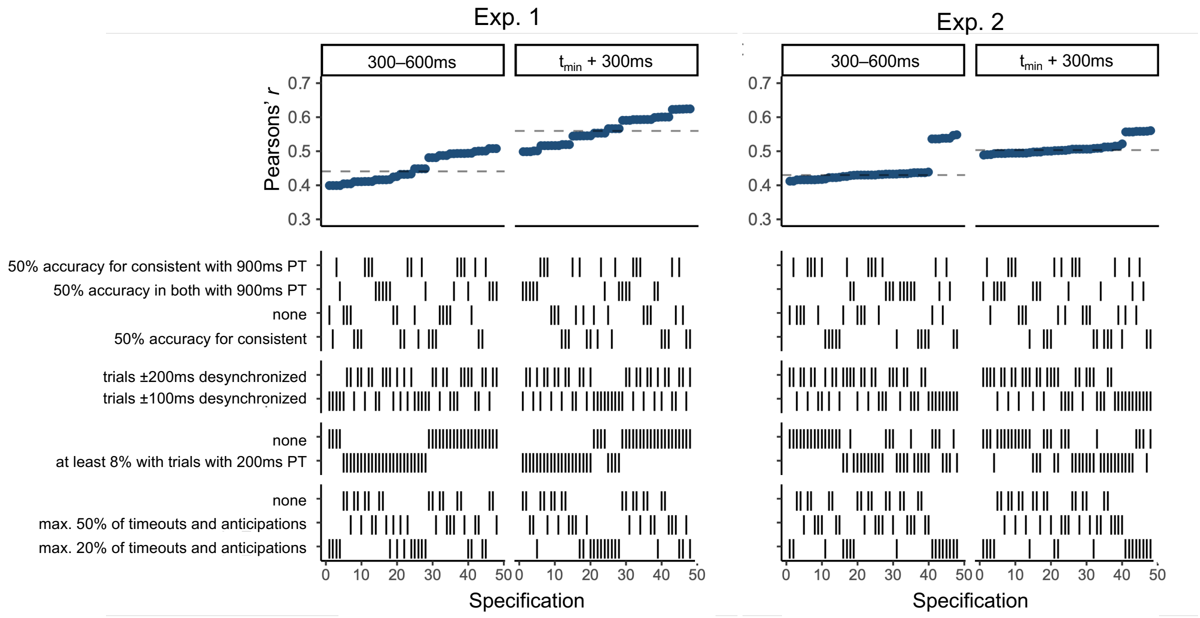


**Supplementary Figure 2: Multiverse analysis for the temporal stability of individual differences in habitual behavior.** We used the set of specifications included in the multiverse analyses conducted in León et al.[1] for the same task. In the top panels, each dot represents a Pearson's r correlation for the proportion of habit errors between test sessions. The facets refer to the preparation times used to calculate the proportion of habitual errors. The dashed line shows the median effect size across specifications. In the bottom panel, combinations of specifications are indicated by vertical lines. PT = Preparation time. Both = Consistent and remapped. Desynchronized = Response desynchronization with the final tone.

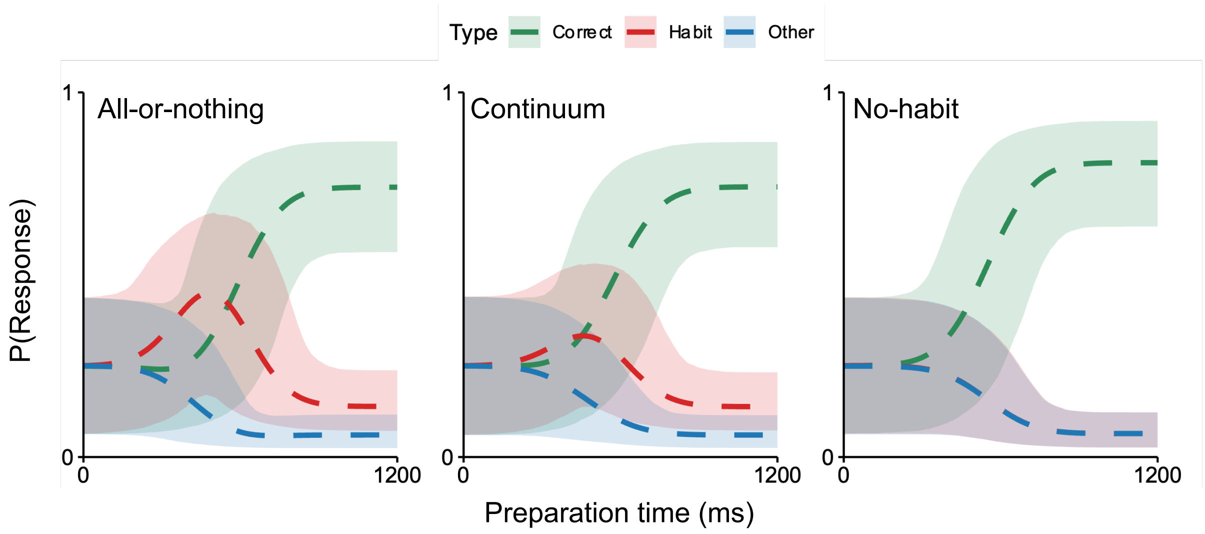


**Supplementary Figure 3: Prior predictive check.** Responses probabilities predicted by the model using the prior distributions specified in the main text. Each dashed line reflects the mean predicted probability, whereas shaded areas reflect the 2.5th and 97.5th percentiles across prior draws. Each group-level parameter was sampled from the prior distributions, and then the response predictions were computed.

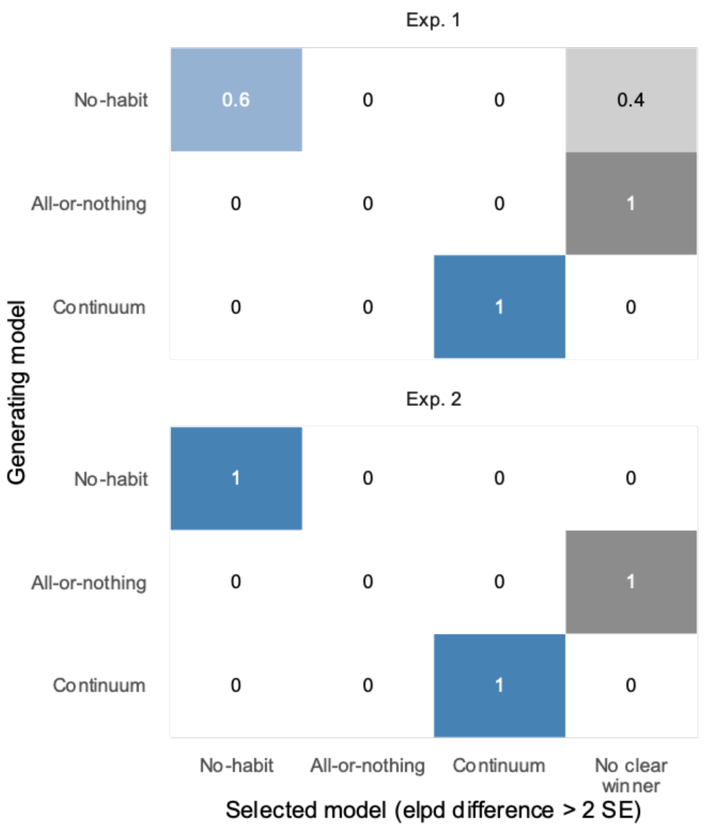


**Supplementary Figure 4: Model recovery.** For each experiment, we simulated behavior from each model 5 times using the posterior-mean parameters of 75 participants in both test sessions, estimated by that model. We fitted each model to each synthetic dataset using the fitting procedure described in the main text as well as the model comparison procedure. A model was considered recovered if its ELPD difference with the second-best model was $> 2 \times$ SE[2]. Otherwise, the dataset was classified as having no clear winner. Each cell indicates the proportion of datasets generated by each model (rows) in which each model was recovered or there was no clear winner (columns). The top row shows results for Exp. 1, the bottom row for Exp. 2. In datasets generated from the all-or-nothing model, there was no clear winner (ELPD difference $< 2 \times$ SE), with a near-tie with the continuum model, because the all-or-nothing model is a special case of the continuum model (i.e., $W_H = 1$). This also occurred in two datasets generated by the no-habit model, which can similarly be considered a special case of the continuum model (i.e., $W_H = 0$ and $\rho = 0$). Importantly, datasets generated from the winning model in our main text (continuum model) were always recovered. Divergent transitions and Pareto k $> 0.7$ never occurred in the fit of the selected model.

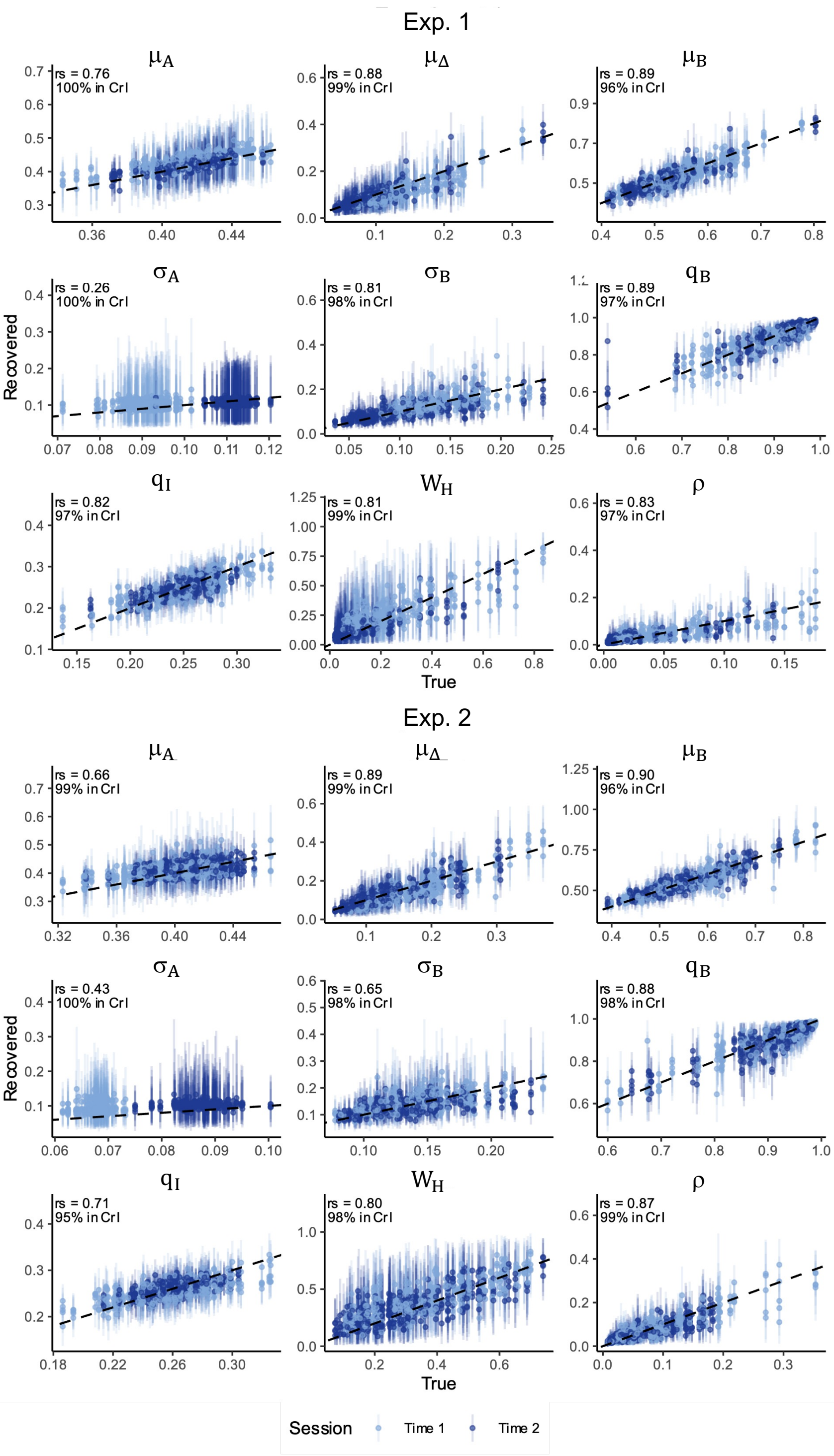


**Supplementary Figure 5: Parameter recovery.** For each experiment, we simulated behavior from the continuum model 5 times using the posterior-mean parameters of 75 participants in both test sessions (150 values), on their own trials. We fitted the model to each synthetic dataset using the fitting procedure described in the main text. Dots indicate posterior means and vertical lines the 95% credible intervals (CrI) of the recovered parameters. Inset text reports Spearman's correlation coefficients (rs) between true and recovered parameters averaged across iterations, and the percentage of 95% CrIs that included the true values. Most parameters were well recovered, with $\geq 95\%$ of CrIs including the true values and point estimates clustering around the "perfect recovery" line (dashed line; ref.[3]). $\mu_A$ showed moderate recovery, while $\sigma_A$ showed poor recovery as reported in prior studies[4,5].

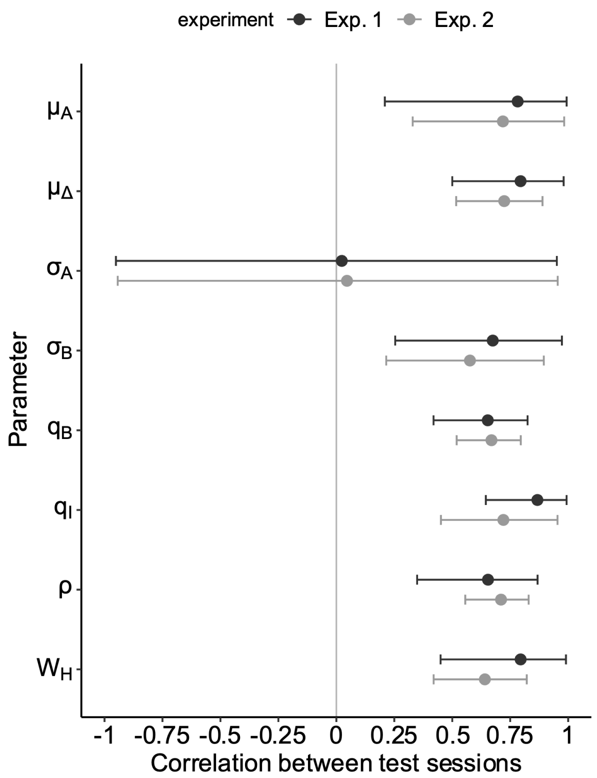


**Supplementary Figure 6: Temporal stability of all the parameters.** Dots illustrate the mean posterior correlation between test sessions of each parameter, and lines reflect the 95 % CrI of the correlation.

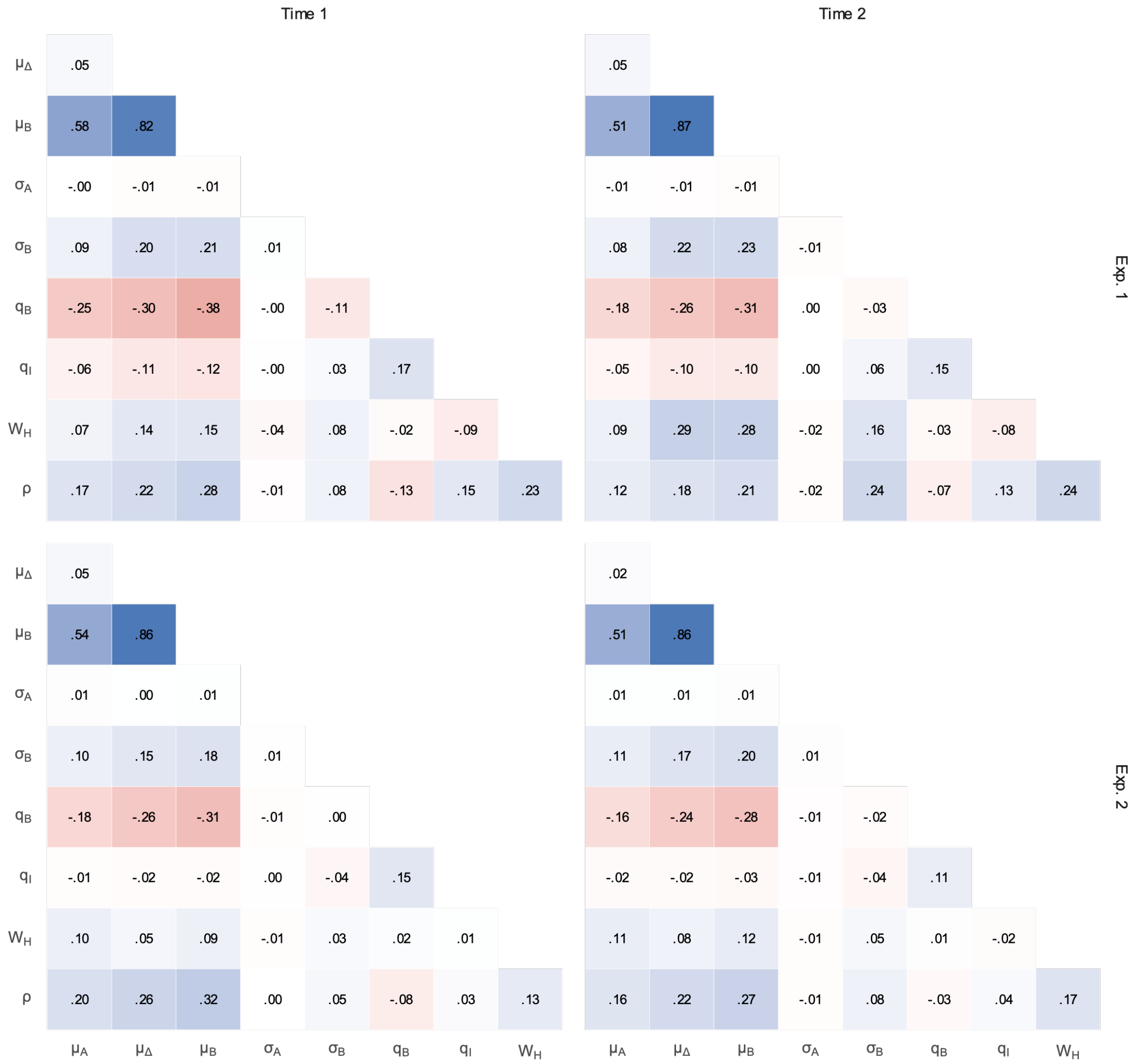


**Supplementary Figure 7: Correlations between participant-level parameters.** Each cell shows the mean posterior of the Pearson correlation between two parameters across participants, separately for each experiment and session. In the model, $\mu_B$ is reparametrized as the sum of $\mu_A$ and $\mu_\Delta$ (see *Methods*), so their correlations with $\mu_B$ are expected.

1 # Supplementary References


2 1. León, J. J., Martínez-López, P., Garre-Frutos, F. & Luque, D. Robust measurement of habits with the forced-response task: A multiverse analysis. (2026) doi:10.31234/osf.io/c4fw7_v2.

3 2. Vehtari, A., Gelman, A. & Gabry, J. Practical Bayesian model evaluation using leave-one-out cross-validation and WAIC. *Statistics and Computing* **27**, 1413–1432 (2017).

4 3. Baribault, B. & Collins, A. G. E. Troubleshooting Bayesian cognitive models. *Psychological Methods* **30**, 128–154 (2025).

5 4. Hardwick, R. M., Forrence, A. D., Krakauer, J. W. & Haith, A. M. Time-dependent competition between goal-directed and habitual response preparation. *Nature Human Behaviour* **3**, 1252–1262 (2019).

6 5. Du, Y. & Haith, A. M. Dissociable habits of response preparation versus response initiation. *Nature Human Behaviour* **9**, 1941–1958 (2025).
